# Netrin1 patterns the dorsal spinal cord through modulation of Bmp signaling

**DOI:** 10.1101/2023.11.02.565384

**Authors:** Sandy Alvarez, Sandeep Gupta, Kaitlyn Honeychurch, Yesica Mercado-Ayon, Riki Kawaguchi, Samantha J. Butler

## Abstract

We have identified an unexpected role for netrin1 as a suppressor of bone morphogenetic protein (Bmp) signaling in the developing dorsal spinal cord. Using a combination of gain- and loss-of-function approaches in chicken, embryonic stem cell (ESC), and mouse models, we have observed that manipulating the level of netrin1 specifically alters the patterning of the Bmp-dependent dorsal interneurons (dIs), dI1-dI3. Altered netrin1 levels also change Bmp signaling activity, as measured by bioinformatics, and monitoring phosophoSmad1/5/8 activation, the canonical intermediate of Bmp signaling, and Id levels, a known Bmp target. Together, these studies support the hypothesis that netrin1 acts from the intermediate spinal cord to regionally confine Bmp signaling to the dorsal spinal cord. Thus, netrin1 has reiterative activities shaping dorsal spinal circuits, first by regulating cell fate decisions and then acting as a guidance cue to direct axon extension.

## Introduction

Netrin1 is a laminin-like protein that was first characterized for its axon guidance activities during embryonic development^1,2^. Netrin1 is widely expressed in both the developing and adult nervous systems, including in the forebrain, optic disc, cerebellum, and spinal cord^1,3–5^, as well as in various tissues outside of the nervous system, including the lung, pancreas, mammary glands, intestine, and developing heart^6–10^.

Many studies have focused on the role of netrin1 directing neural circuit formation in the developing spinal cord, where it was initially identified^1,2^. Spinal neurons arise at stereotyped positions along the dorsal-ventral axis, such that the dorsal spinal cord is comprised of at least six populations of dorsal interneurons (dIs), dI1-dI6, that are derived from six dorsal progenitor (dP) domains, dP1-dP6^11,12^. This pattern is generated by multiple signals, which collectively act on proliferating neural progenitor cells (NPCs) in the ventricular zone (VZ)^11,12^. In the dorsal spinal cord, these signals include multiple members of the bone morphogenetic protein (Bmp) family, which are secreted from the roof plate (RP) at the dorsal midline of the spinal cord^13,14^. Bmps are sufficient for the specification of the RP itself, as well as the dI1s, dI2s, and dI3s^15–19^. The Bmps activate distinct type I Bmp receptors^18,20^, which in turn phosphorylate the receptor-regulated (R)-Smads, Smad1/5/8, the intracellular mediators of Bmp signaling^21^, to direct dPs to differentiate into post-mitotic dIs^22^. Bmps have reiterative roles, directing both NPC proliferation and dI differentiation^18^. Additional signals include retinoic acid (RA), which is present in the surrounding paraxial mesoderm and is important for patterning and neuralization^23–25^. Immediately after neurogenesis, dIs start to extend axons toward their synaptic targets^26^. Netrin1 acts to direct dI1 (commissural) axons towards the floor plate (FP), at the ventral midline of the spinal cord^1^. While *netrin1* is expressed by both NPCs in the intermediate VZ, and the FP in the mouse spinal cord, recent studies have suggested that it is the NPC-derived netrin1 that is most critical for mediating axon guidance events^27–30^. NPC-derived netrin1 is thought to be trafficked along the radial processes of the NPCs until it is deposited on the pial surface, where it forms an adhesive growth substrate that locally orients dI1 axon extension^28–30^.

The netrin family has subsequently been shown to play many critical roles in developmental and physiological processes beyond axon guidance. Netrin1 is involved in the progression of cancers^31–35^, diabetes^36^, and inflammatory bowel diseases^37^. Netrin1 also directs cellular differentiation across organ systems. In the skeletal system, netrin1 mediates bone remodeling by suppressing osteoclast differentiation and promoting osteoblast differentiation^38^. Netrin1 also plays a role in the morphogenesis and differentiation of the mouse mammary glands^39^ and can induce human embryonal carcinoma cells to differentiate into a neuroectodermal-like cell type^40^. However, no role for netrin1 directing cell fate in the developing nervous system *in vivo* has been described.

Here, we provide the first evidence that netrin1 can regulate cell fate specification in the dorsal spinal cord. Since netrin1 has been previously shown to suppress the Bmp signaling pathway in different cell types *in vitro*^41^, we sought to investigate the relationship between netrin1 and Bmp signaling in the patterning of the embryonic spinal cord. These studies support the hypothesis that netrin1 acts from the intermediate spinal cord to regionally limit the extent of Bmp signaling to the dorsal spinal cord. Using a combination of bioinformatics with gain and loss of function approaches in chicken, mouse and stem cell models, we have found that modulating the level of netrin1 has profound effects on the number of the Bmp-dependent dIs, i.e., dI1-dI3s. Netrin1 appears to mediate its effects through the Bmp pathway, given that changes in dI number were accompanied by alterations in the levels of both phosopho (p) Smad1/5/8 and inhibitor of differentiation/DNA-binding (Id) expression. The Id family are key downstream mediators of Bmp signaling^42^, that modulate the activities of the proneural basic helix-loop-helix (bHLH) proteins^43–45^ to prevent exit from the cell cycle^46,47^. Thus, activation of Ids can result in NPCs being held in a proliferative state.

Together, these findings suggest a new role for netrin1 in the developing spinal cord, modulating Bmp signaling to fine-tune neural patterning. Thus, netrin1 has an earlier role than previously realized, with reiterative activities shaping dorsal spinal circuits, first by regulating cell fate decisions and then acting as a guidance cue to direct axon extension. Both netrin1 and members of the Bmp family are widely expressed, with the Bmps also having reiterative roles in cell growth, differentiation, migration, apoptosis, and homeostasis in the developing embryo and adult^48–50^. These studies therefore also open the possibility that netrin1 can modulate Bmp-dependent processes in other organs.

## Results

### Netrin family expression in the developing spinal cord

Netrin1 has widespread expression in the developing mouse spinal cord; it is expressed by cells in the FP and the ventral and intermediate NPCs, resulting in netrin1 protein decorating the ventral and intermediate pial surface^29,30^. In contrast, netrin function is mediated by both netrin1 and netrin2 in the embryonic chicken spinal cord, which have distinct distributions at different stages (Figure 1A-1L)^1,51^. At Hamburger Hamilton (HH) stage 18, when dI fate specification commences^18^, netrin1 is present in the ventral spinal cord (Figure 1A, 1B), while netrin2 is present in the intermediate spinal cord, and is absent from the dorsal- and ventral-most regions (arrowheads, Figure 1F). By HH stage 24, when axonogenesis is ongoing, *netrin1* expression is confined to the FP (arrowhead, Figure 1I), while the domain of netrin2 has contracted to an intermediate region that spans from immediately below the dorsal root entry zone to just above the motor column (arrowheads, Figure 1K, L). In both cases, the distribution of *netrin* mRNA is largely distinct from the distribution of netrin protein. This is most evident for netrin2: *netrin2* mRNA is present in the ventricular zone (VZ), while netrin2 protein accumulates on the pial surface immediately adjacent to its expression domain. Thus, the expression patterns of chicken netrin1 and netrin2 are the composite of the mouse netrin1 distribution, including the conserved upper boundary in the dorsal spinal cord.

**Figure 1.**
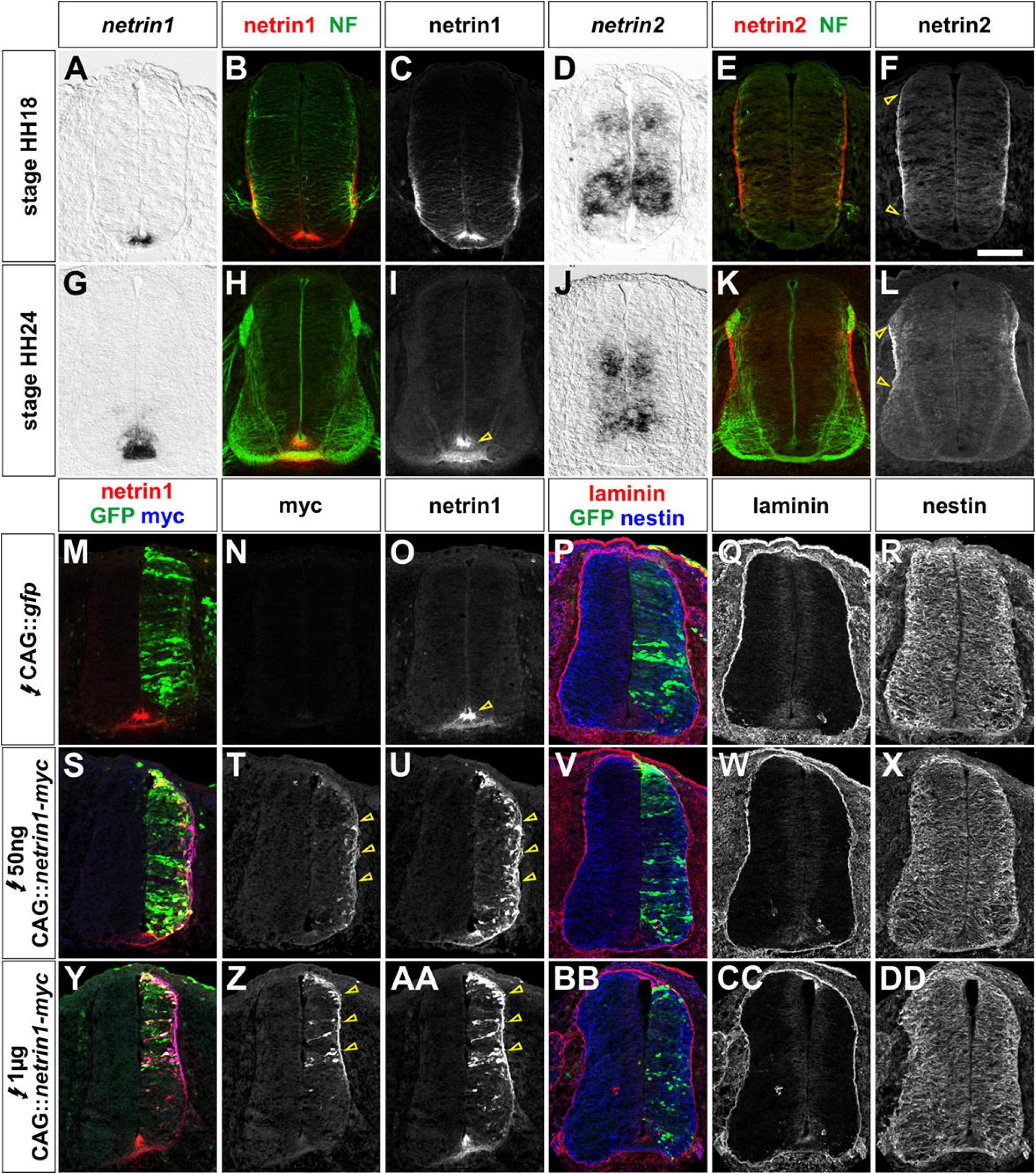
Overexpression of netrin1 does not affect the integrity of the developing spinal cord (A-L) Distribution of netrin1 (A-C, G-I) and netrin2 in (D-F, J-L) in thoracic sections of the HH18 (A-F) and HH24 (G-L) spinal cord. *Netrin1* mRNA is expressed in the apical FP (A, G), while netrin1 protein (red, B, H) decorates the apical-most and basal FP (arrowhead, C, I), where it is co-incident with the NF^+^ axons crossing the FP (H). *Netrin2* mRNA is expressed in the intermediate VZ (D, J), while netrin2 protein (red, E, K) decorates pial surface in the intermediate spinal cord (arrowheads, F, L), immediately adjacent to NF^+^ axons (green) extending ventrally in the dorsal spinal cord (L). (M-R) Electroporation of a control fluorophore, GFP (green, M, P), expressed from a ubiquitously expressed CAG enhancer, does not affect the distribution of endogenous netrin1 (red, M; arrowhead, O), or the integrity of the spinal cord as assessed by antibodies against laminin (red, P, Q) and nestin (blue, P, R). (S-DD) In contrast, electroporation of a low (50ng, S-X) or high (1μg, Y-DD) concentration of netrin1-myc construct, results in the ectopic netrin1 (red, S, Y, U, AA) and myc (blue, S, Y, T, Z) decorating the pial surface of the spinal cord (arrowheads, T, U, Z, AA). However, there is no effect on the distribution of either laminin (red, V, W, BB, CC) or nestin (blue, V, X, BB, DD). Scale bar: 100µm

### Netrin1 misexpression does not perturb the architecture of the spinal cord

To investigate whether netrin1 has effects in the spinal cord distinct from its role mediating axon guidance, we first assessed whether the misexpression of netrin1 altered the general structure of the chicken spinal cord. A range of concentrations (50ng, 500ng, 1μg) of C-terminally myc-tagged netrin1 (netrin1-myc) were electroporated into the HH stage 14 spinal cord under the control of the ubiquitously expressed CAG enhancer^52^, and the consequences examined two days later, at HH stage 24/25. CAG::*gfp* was concomitantly electroporated in all experiments, to both serve as a control (Figure 1M-1R), and indicate the extent of electroporation. Initially, both chicken and mouse netrin1 were used in these experiments, which are ∼90% similar at the amino acid level. However, since their activities were found to be very similar (data not shown), we proceeded with mouse netrin1, which could be additionally identified as a distinct signal using species specific antibodies. While the GFP fluorophore had no effect on the distribution of endogenous chicken netrin1 (arrow, Figure 1M, 1O), the misexpression of netrin1-myc resulted in both myc and netrin1 being targeted to the pial surface (arrows, Figure 1S-1U, 1Y-1AA). Increasing the concentration of electroporated netrin1-myc increased the amount of pial-myc (compare arrows, Figure 1T and 1Z). However, even at the highest levels of exogenous netrin1-myc, there was no effect on the integrity of the laminin^+^ basal membrane (Figure 1V, 1W, 1BB, 1CC), or the nestin1^+^ radial processes of the neural progenitor cells (NPCs; Figure 1X, 1DD), compared to control electroporations (Figure 1P-1R). Thus, misexpression of netrin1 does not change the overall architecture of the spinal cord.

### Netrin1 overexpression in chicken embryos results in a dose dependent reduction in dIs

Although the general architecture of the spinal cord was unaffected, we did observe a reduction in the size of the spinal cord after electroporation with netrin1-myc. To further assess this phenotype, we quantified the area bounded by either the Sox2^+^ NPCs or p27^+^ post mitotic neurons in control (Figure 2A-2C, 2N) versus netrin1-myc (Figure 2D-2F, 2N) electroporations. In the control condition, the electroporated vs. non-electroporated sides of the embryo were statistically indistinguishable for both Sox2^+^ NPCs (p>0.65, Student’s *t*-test) or p27^+^ neurons (p>0.60). In contrast, there was a ∼25% reduction in the total area bounded by Sox2^+^ NPCs and ∼33% decrease in area of the p27^+^ neurons after ectopic expression of netrin1-myc (Figure 2N). This reduction was seen with all concentrations of netrin1, suggesting that netrin1 can potently affect the number of neurons that arise in the spinal cord.

**Figure 2.**
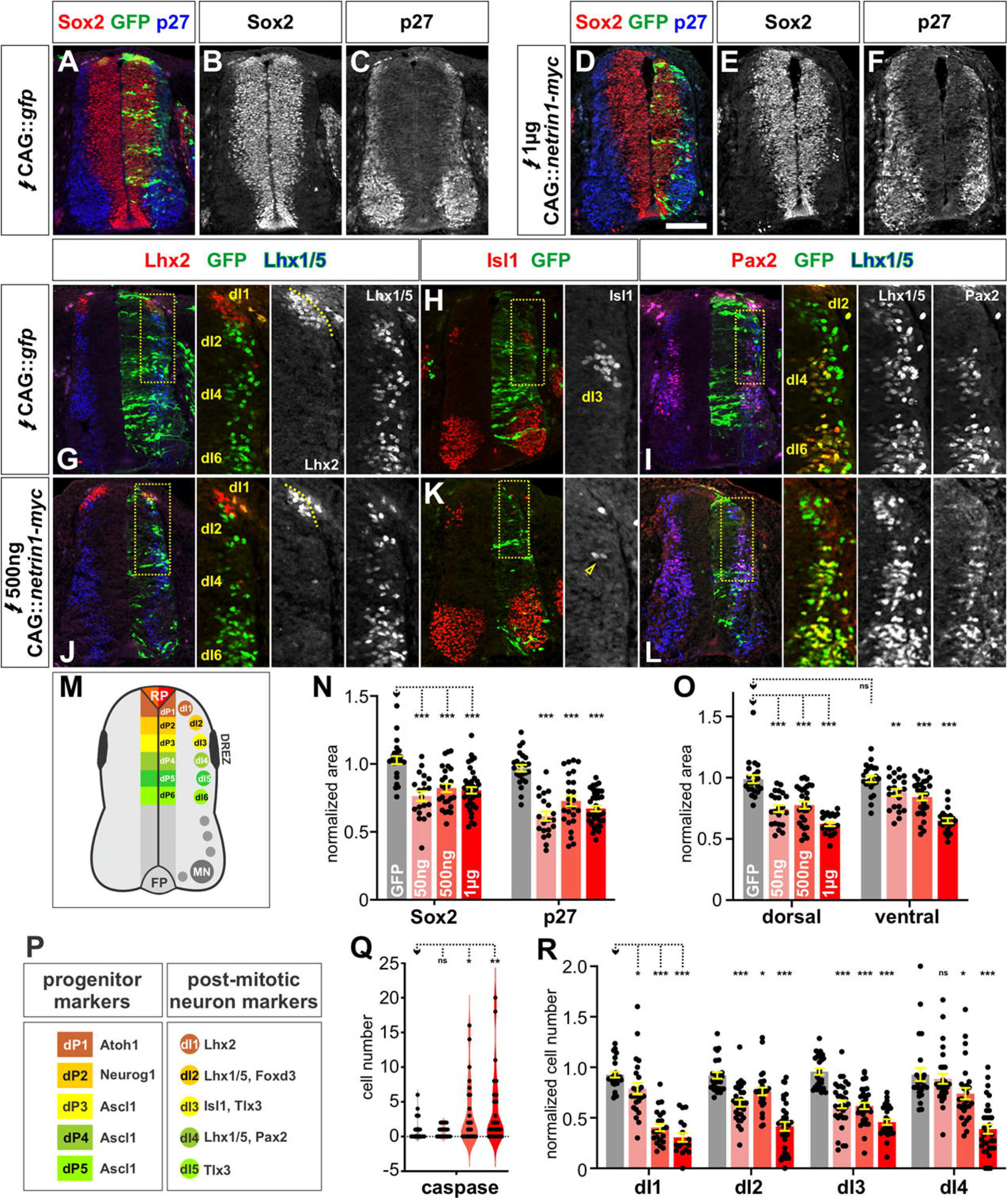
Overexpression of netrin1 in chicken embryos results in the loss of dorsal interneurons (A-L) Chicken spinal cords were electroporated at HH stage 14 with *Gfp* (A-C, G-I) or different concentrations of *netrin1* (50ng, 500ng, 1μg) (D-F, J-L) under the control of the CAG enhancer, and incubated until HH stage 24. Thoracic transverse sections were labeled with antibodies against Sox2 (red, A, B, D, E), p27 (blue, A, C, D, F), Lhx2 (red, G, J), Isl (red, H, K), Lhx1/5 (blue/green G, I, J, L) and Pax2 (red, I, L). The dotted box (G-L) indicated the magnified region in the adjacent panel(s). (M) Schematic transverse section of the spinal cord, showing the position of the dorsal progenitor (dP) domains, and post-mitotic dorsal interneurons (dIs). (N, O) Ectopic *Gfp* expression had no significant effect on the general specification of the spinal cord. In contrast the total area occupied by Sox2^+^ (progenitors) or p27^+^ (neurons) cells was significantly reduced at all concentrations of netrin1 tested (N). The dorsal spinal cord was more profoundly affected at lower concentrations of netrin1 than the ventral spinal cord (O). A comparison of the dorsal vs. ventral area of n= >20 sections, from 4 embryos (GFP, 50ng, 500ng, 1μg netrin), Student’s *t*-test. (P) The different classes of dIs can be identified by specific combinations of transcription factors. (Q) There was no significant difference in the number of cells were caspase^+^ (H, p>0.58) between spinal cords that were electroporated with GFP and 50ng netrin1. In contrast, there were significant differences when the higher concentrations of netrin1 where electroporated. n= >20 sections, from 4 embryos (GFP, 50ng, 500ng, 1μg netrin). (R) Ectopic *Gfp* expression also had no significant effect on dI specification. However, all concentrations of netrin1 tested were able to significantly reduce the number of Lhx2^+^ dI1s, Lhx1/5^+^ Pax2^−^ dI2s and Isl1^+^ dI3s in a dose dependent manner. In contrast, only the higher concentrations of netrin1 reduced the number of Lhx1/5^+^ Pax2^+^ dI4s. n= >20 sections, from 6 embryos (GFP, 50ng, 500ng, 1μg netrin), One way ANOVA. Probability of similarity between control and experimental groups: *= p < 0.05, **p<0.005 *** p<0.0005. Scale bar: 100µm

While the size of the entire spinal cord was reduced, the effect of netrin1 on the dorsal spinal cord was more pronounced and observed at lower concentrations (Figure 2O). To further assess the consequence of netrin1 misexpression on specific dorsal identities, we used a well described panel of antibodies against transcription factors that distinguish specific dI fates^53^, to monitor the numbers of Lhx2^+^ dI1s, Lhx1/5^+^ Pax2^−^ dI2s, Isl1^+^ dI3 and Lhx1/5^+^ Pax2^+^ dI4s. We observed that the Bmp-dependent populations^25^, i.e., dI1, dI2 and dI3, are all significantly reduced after netrin1-myc electroporation compared to the GFP control. These reductions were concentration-dependent, such that more dI1/dI2/dI3s were lost, as the amount of netrin1-myc increased. ∼75% of dI1s and ∼50% of dI2/dI3s were ablated at the highest concentration of netrin1-myc tested (Figure 2RR). We also observed a profound loss of dI1 commissural axons in the remaining dI1 population (Figure S2).

In contrast, the dI4s, a Bmp-independent population, were less profoundly affected. The lowest level of netrin1-myc did not significantly affect the numbers of dI4s, rather dI4s were only lost as the concentration of netrin1-myc increased (Figure 2R). The more widespread loss of neurons resulting from the highest levels of netrin1-myc misexpression may be a consequence of cell death. The number of caspase^+^ cells was not significantly different between control and 50ng CAG::netrin1-myc, but did increase as the concentration of netrin1-myc increased (Figure 2Q).

### Addition of netrin1 to mESCs blocks their ability to acquire dI1/dI3 fates

We next assessed the activity of netrin1 in stem cell model that recapitulates the early events that direct cell fate in the developing spinal cord^25^. In brief, bFGF/Wnt signaling directs mouse embryonic stem cells^54^ (mESCs) into a bipotential neuromesodermal progenitor (NMP) fate, that is a critical intermediate for the cells of the caudal neural tube^55^ (Figure 3A). Our recent work has shown that addition of retinoic acid (RA), from day 3-5, directs NMPs to a caudal dorsal progenitor (dP) fate, specifically that of the intermediate neural tube, ultimately resulting in the specification of dI4, dI5 and dI6^25,56^. The sequential addition of Bmp4 from day 4-5 further dorsalizes the NMPs into the dPs that specify the dI1, dI2 and dI3 fates. Thus, these RA±Bmp4 directed differentiation protocols provide an additional model to investigate the mechanisms that drive dorsal spinal cord development.

**Figure 3:**
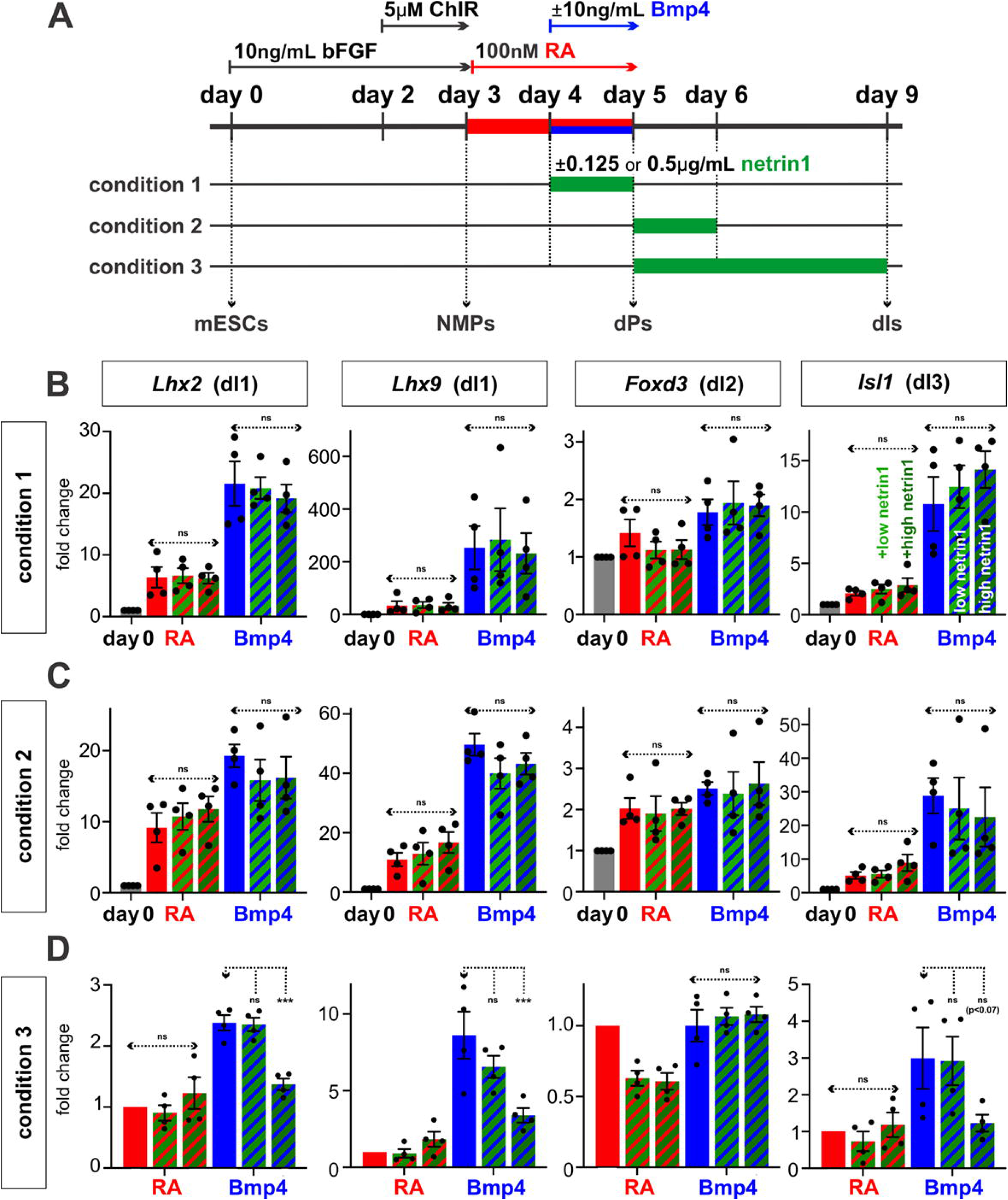
Addition of netrin1 blocks dorsalization in mESC stem cell model of dI differentiation. (A) Two concentrations of Netrin1 recombinant protein (0.125μg/ml (low) and 0.5μg/ml (high)) were added to the RA±Bmp4 protocol at three different timepoints: alongside with Bmp4 from day 4-5 (condition 1); immediately after Bmp4 treatment from day 5-6 (condition 2), and for an extended period after Bmp4 treatment from day 5-9 (condition 3). qPCR was used to assess alterations in gene expression. (B, C) The addition of netrin1 in condition 1 and 2 had no significant effect on the expression of *Lhx2* and *Lhx9* (dI1), *Foxd3* (dI2) or *Isl1* (dI3) compared to RA or RA+Bmp4 controls. (D) Prolonged treatment with 0.5μg/ml netrin1 in the RA+Bmp4 protocol significantly reduced the expression of the dI1 markers, and there is a trend (p<0.07) towards the loss of dI3 marker *Isl1*. Probability of similarity between control and experimental groups: *= p < 0.05

Using genomic data from our prior studies^25,57^ we determined that *netrin1* is expressed by mESC-derived progenitors, but generally absent from differentiating neurons (Figure S1A), as *in vivo*^2,30^. We assessed the effect of adding two concentrations of exogenous netrin1 - 0.125 μg/ml (low) and 0.5 μg/ml (high) - to mESC-derived NMPs in the RA±Bmp4 protocols at three different timepoints. Netrin1 was added concomitantly with Bmp4 from day 4-5 (condition 1); immediately after Bmp4 treatment from day 5-6 (condition 2); and finally, for an extended period after Bmp4 treatment from day 5-9 (condition 3) (Figure 3A). The cultures were then assessed for the specification of the dorsal-most dIs at day 9, using qPCR analyses (Figure 3B-3D). The addition of netrin1 with, or immediately after, Bmp4 treatment had no apparent effect on the identity of the cultures (Figure 3B, 3C). We also found no effect on *Foxd3* expression in any of the conditions, i.e., dI2s continue to assume their fate in the presence of netrin1. However, prolonged treatment with 0.5 μg/ml netrin1 in the RA+Bmp4 protocol significantly reduced the expression of *Lhx2* and *Lhx9*, both markers of dI1s, and there is a trend (p<0.07) towards the loss of *Isl1*, a dI3 marker. Thus, the extended treatment of stem cell derived NMPs with high levels of netrin1 is sufficient to prevent some dorsalization. This result, coupled with the observation that netrin1 misexpression in the chicken spinal cord most effectively suppresses the Bmp-dependent dIs, suggests that netrin1 is sufficient to counteract the activities of the Bmps.

### The loss of netrin1 increases the size and number of the dorsal most spinal progenitors

We next assessed the requirement for netrin signaling on dorsal spinal fate specification using mice deficient for *netrin1*^29,58,59^. We analyzed a null allele for netrin1 (*ntn1^−/−^*) which was previously analyzed for axon guidance defects in the developing spinal cord^60^, but was not evaluated for changes in dorsal cell fate. We focused our analysis on embryonic (E) day 11.5, when dorsal fate specification is robustly ongoing in the spinal cord. We first observed a marked expansion of the dorsal most progenitor domains flanking the RP in the *netrin1* mutants. The dP1 domain is almost 2-fold larger in size (p<0.0001 significantly different compared to littermate control) and there is a ∼25% increase in the number of Atoh1^+^ dP1s (p<0.045; Figure 4A, 4D, 4M). The dP2 domain is ∼60% larger (p< p<0.0001; Figure 4A, 4D, 4M), as measured by the area bounded by the bottom of the Atoh1^+^ dP1 domain and the top of Ascl1^+^ dP3 domain. There is also a ∼40% increase in the area (p<0.0001) demarked by Ascl1^+^ cells, which form the dP3-dP5 domain (Figure 4A, 4D, 4M). In contrast, we observed no significant difference in the number of Olig2^+^ cells in control and mutant sections (p>0.25; Figure 4B, 4E, 4M), suggesting that there was no effect on the motor neuron progenitor domain (pMN)^61^. Together, these results suggest that the loss of *netrin1* increases the number/size of the most dorsal neural progenitors in the spinal cord. The increase in dPs did not stem from an increase in the rate of cell division, since there was no significant change in the number of pH3^+^ cells in mitosis (p>0.5; Figure 4C, 4F, 4G), or from altered patterns of cell death (p>0.055; Figure 4H).

**Figure 4.**
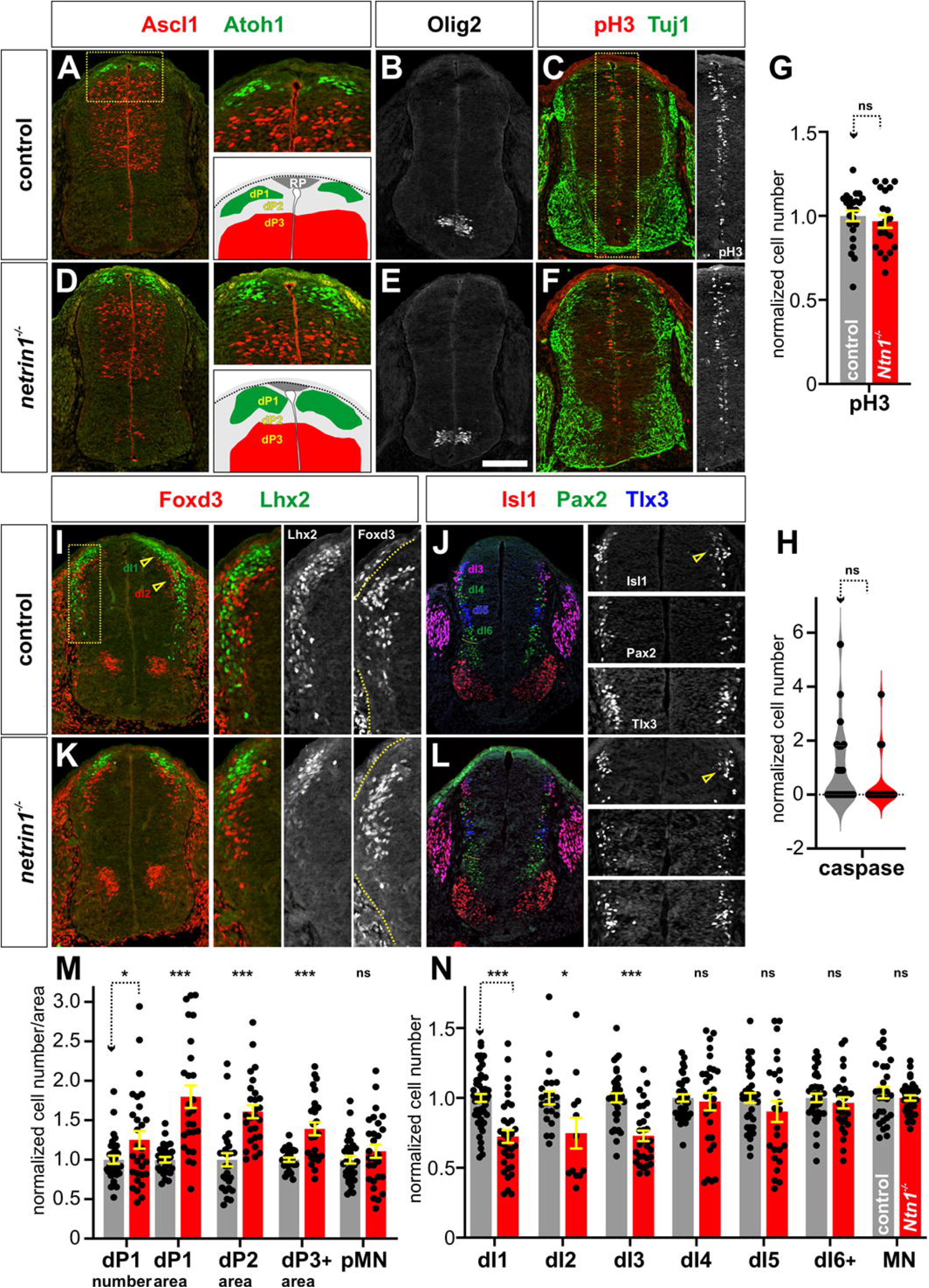
Netrin1 is required to limit the number of the dorsal most NPCs (A-F) Thoracic transverse spinal cord sections from either control (A-C) or *netrin1^−/−^* (D-F) E11.5 mouse spinal cords were labeled with antibodies against Ascl1 (A, D; red; dP3-dP5), Atohl1 (A, D; green, dP1), Olig2 (B, E; pMN), phospho-histoneH3 (C, F; red; cells in mitosis (M) phase), and Tuj1 (C, F; green; neurites). The dotted box in A, C indicates position of the magnified region in the adjacent panel(s). (G, H) There was no significant difference in the number of cells either in M-phase (G, p>0.50; n= > 20 sections, from 3 control and 3 *netrin1^−/−^*embryos) or that were caspase^+^ i.e. dying (H, p>0.28; n= >13 sections, from 3 control and 3 *netrin1^−/−^*embryos) between control and *netrin1^−/−^* spinal cords. (I-L) To assess for changes in the number of post mitotic dIs, thoracic transverse spinal cord sections from either control (I, J) or *netrin1^−/−^*(K, L) E11.5 mouse spinal cords were labeled with antibodies against Lhx2 (I, K; green; dI1), Foxd3 (I, K; red; dI2), Isl (J, L; red, dI3, MNs), Pax2 (J, L; green; dI4, dI6, v0), and Tlx3 (J, L; blue; dI3, dI5). The dotted box (I, K, J, L) indicates the magnified region in the adjacent panel(s). (M) Loss of netrin1 resulted in a 25% increase in the number of Atoh1^+^ dP1s and an almost 2-fold increase in the area occupied by the dP1s. Similarly, the area of the dP2 domain (region bounded by the Atoh1^+^ and Ascl1^+^ domains) was increased by 60%, and the Ascl1^+^ dP3-dP5 domain was increased by 40% (n= > 26 sections, from 4 embryos). (N) This increased number of progenitors did not result in a loss of dIs. Rather there was a ∼30% decrease specifically in the number of dI1, dI2 and dI3s (n= > 25 sections, from 5 embryos), but not in the intermediate dorsal populations or the ventral motor neurons (MNs), Probability of similarity between control and experimental groups: *= p < 0.05, ** p<0.005, *** p<0.0005; Student’s *t*-test. Scale bar: 100µm

We next assessed whether the increased number of dorsal progenitors affected the number of post-mitotic dIs. To our surprise, we found that there was a ∼30% decrease in the number of Lhx2^+^ dI1s (p<0.0001; Figure 4I, 4K 4N), Foxd3^+^ dI2s (p<0.017; Figure 4I, 4K, 4N) and Isl1^+^ Tlx3^+^ dI3s (p<0.0001) (Figure 4J, 4L, 4N). In contrast, there was no significant difference in the more intermediate dIs, i.e., the Pax2^+^ dI4s (p>0.67), Tlx3^+^ Isl1^−^ dI5s (p>0.19), and Pax2^+^ dI6s/v0/v1 (dI6+; p>0.48), or the Isl1^+^ MNs (p>0.36). Thus, the loss of netrin1 appears to specifically affect the development of the Bmp-dependent dorsal most dIs and does not affect the Bmp-independent intermediate dIs, and ventral spinal cord. This result is consistent with the hypothesis that netrin1 regulates the activity of Bmps in the dorsal spinal cord.

### Netrin1 regulates Bmp signaling in stem cell-derived dorsal progenitors

To further understand the mechanism by which netrin1 regulates dI specification, we assessed the transcriptomic profiles of mESC-derived neural progenitors undergoing the RA±Bmp4 directed differentiation protocol in the presence of netrin1 protein (Figure 3A). Samples for bulk RNA-seq analyses were collected on day 5 (condition 1), day 6 (condition 2) and day 9 (condition 3) (Figure 5A). Surprisingly, we observed almost no transcriptional changes between the control and netrin 1 treated samples in condition 1 (1 gene down regulated, FDR, p<0.05). In contrast, there were a modest number of differentially expressed genes in condition 2 (26 downregulated, 19 upregulated, FDR p<0.05) and a substantial number in condition 3 (4847 downregulated, 4125 upregulated, FDR p<0.05) (Figure 5B, 5C). Thus, netrin1-mediated transcriptional changes only occur once progenitors commit to a dorsal identity.

**Figure 5:**
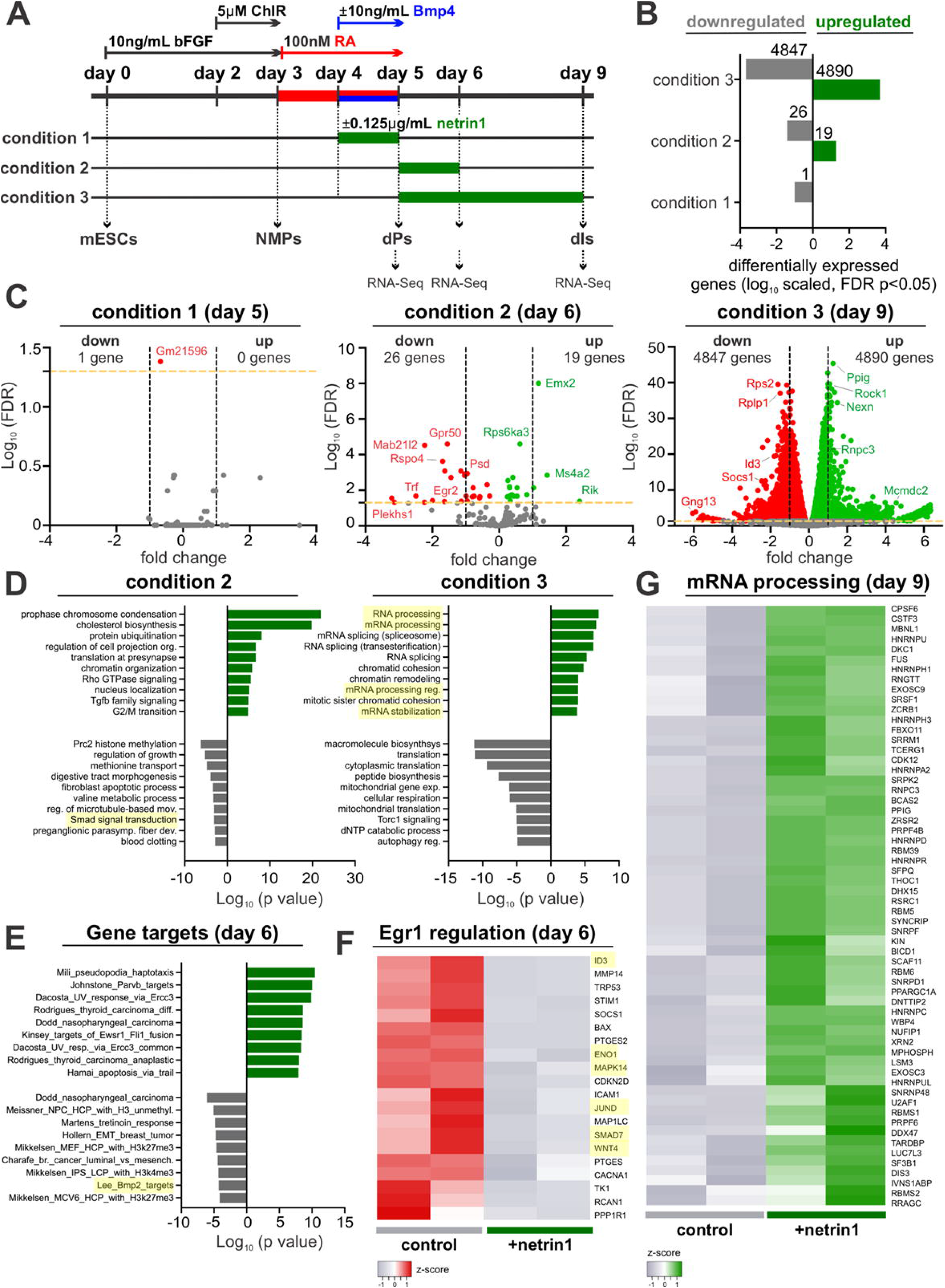
Netrin1 downregulates BMP signaling and alters mRNA processing (A) Netrin1 was added to the RA±Bmp4 directed differentiation protocol at three different timepoints. RNA samples for bulk RNA-Seq were collected on day 5 (condition 1), day 6 (condition 2) and day 9 (condition 3). (B, C) The pulse of netrin1 in condition 1 resulted in essentially no differentially expressed genes at a false discovery rate (FDR) of p<0.05. In contrast, there was a modest increase in transcriptional changes in condition 2, while ∼10,000 genes were differentially expressed in condition 3, after extended treatment with netrin1. (D) Gene ontology analyses of differentially expressed genes due to netrin1 treatment, showed that Bmp signaling was downregulated at day 6 and mRNA processing was upregulated at day 9. (E) A gene target analysis of published gene sets, also identifies that Bmp2 target genes (highlighted) are downregulated after netrin1 addition by day 6. (F) A number of transcription factor regulatory networks (for complete set of networks, see Table S1) where identified as being downregulated in netrin1 treated cultures in condition 3. Many BMP target genes were found to be downregulated by Egr1 (highlighted), including Id3. The heatmap shows the FPKM values of select genes. (G) Heatmap showing the upregulated expression (FPKM values) of mRNA processing genes in netrin1 treated cultures at day 9 (condition 3), validating the GO analysis in panel D.

To gain insight into the signaling pathways affected by netrin1 treatment, we conducted Gene Ontology analyses of condition 2 and 3 (Figure 5D). Supporting the hypothesis that netrin1 suppresses the Bmp signaling pathway, the Smad signal transduction gene module was downregulated at day 6 (Figure 5E). Similarly, we observed that Bmp2 target genes^62^ were among the gene sets associated with previously published studies that were downregulated in condition 3. In particular, the gene network regulated by the Egr1 transcription factor was downregulated in condition 3, which includes many known transcriptional targets of Bmp signaling, such as *Id3*, *Smad7*, *Wnt4*, and *Mapk14* (Figure 5F, Table S1). Egr1 has also been described as being regulated by Bmp signaling^63^. By day 9, the upregulated GO signatures suggest that netrin1 regulates mRNA processing (Figure 5D). Many upregulated genes in condition 3 are associated with mRNA processing and splicing (Figure 5G), such *Ppig*, a RNA binding protein^64^, and *Rnpc3*^65^, which encodes part of the spliceosome. Taken together, these transcriptomic analyses suggest a mechanism where netrin1 acts to inhibit Bmp signaling, perhaps by regulating the mRNA processing of Bmp target genes, such as the Ids, to control dI fate specification.

### The gain or loss of netrin1 activity alters the level of Bmp signaling

We next directly examined whether modulating netrin1 levels affects Bmp signaling, by monitoring phosphorylated (p) Smad1/5/8 levels^66,67^ in Cos7 cells *in vitro.* Cos7 cells can endogenously transduce Bmp signaling; the level of pSmad1/5/8 robustly increases when Cos7 cells are treated with Bmp4 for an hour (Figure 6H, 6K). However, if 0.5µg/mL netrin1 is added concomitantly with Bmp4, pSmad1/5/8 levels decrease by ∼60%. This is a dose-dependent response: halving the amount of netrin1 diminishes this response, while 0.125µg/mL netrin1 treatment has no significant effect on Smad1/5/8 activation (Figure 6H, 6K).

**Figure 6.**
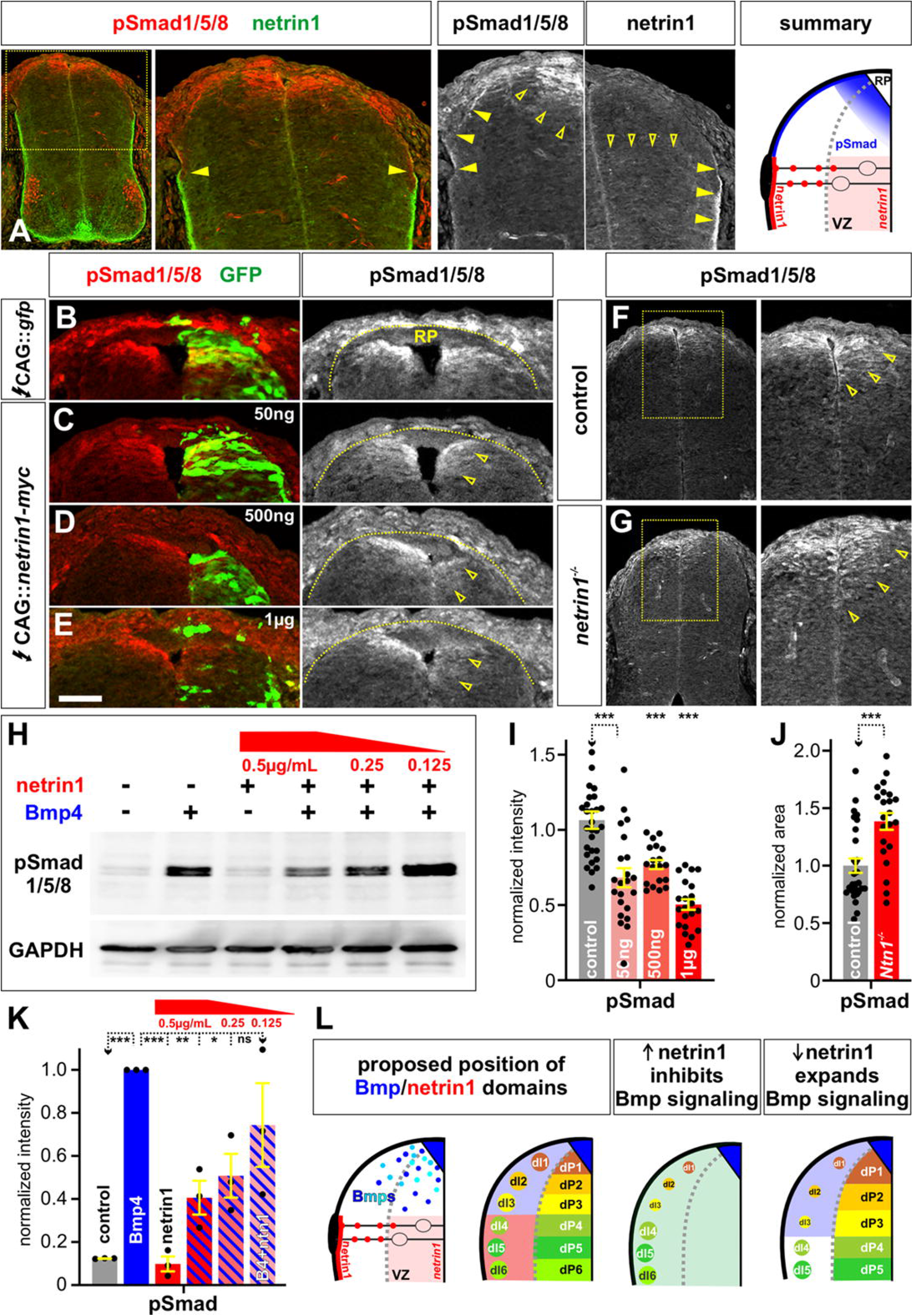
Netrin1 modulates the level of Bmp signaling both *in vivo* and *in vitro* (A) Thoracic sections of E11.5 mouse spinal cord labeled with antibodies against pSmad1/5/8 (red) and netrin1 (green). The pSmad1/5/8 and netrin1 proteins are detected in neighboring domains. pSmad1/5/8 is present in the VZ immediately flanking the RP (open arrowheads), and along the pial surface of the dorsal most spinal cord (closed arrowheads). Netrin1 is present in the intermediate spinal cord, at low levels in the VZ (open arrowheads) where *netrin1* is expressed, and high levels on the pial surface (closed arrowheads) after trafficking along the radial processes (summary)^29^. (B-E, I) Chicken spinal cords were electroporated at HH stage 14 with *Gfp* (A) or different concentrations of *netrin1* (50ng, 500ng, 1µg) (B-D) under the control of the CAG enhancer, and incubated until HH stage 24/25. Thoracic transverse sections were labeled with antibodies against pSmad1/5/8 (red). Ectopic netrin1 in the dorsal most spinal cord resulted a ∼30-50% decrease in the levels of Smad1/5/8 compared to a control GFP electroporation. (F, G, J) Thoracic transverse spinal cord sections from either control (F) or *netrin1^−/−^*(G) E11.5 mouse spinal cords were labeled with antibodies against pSmad1/5/8. The loss of netrin1 resulted in a ∼40% larger Smad^+^ area (J), suggesting Bmp signaling had been increased. (H, K) The interaction between netrin1 and Bmp4 was further assessed in a western analysis, using GAPDH levels as a loading control. Treating Cos7 cells with Bmp4 resulted in the robust activation of pSmad1/5/8; while treatment with netrin1 alone had no effect on Smad activation above control levels. However, if netrin1 is added together with Bmp4, there is a decrease in Smad activation in a dose-dependent manner. The highest level of netrin1 (0.5ng/mL) resulted in a ∼60% decrease in the level of pSmad1/5/8, suggesting that Bmp signaling had been suppressed. (L) Model for the biological significance of the netrin1/Bmp interaction. Multiple Bmps are secreted from the roof plate where they act to pattern the surrounding tissue into the dorsal progenitor domains (dP1-dP3). Netrin1 acts as a boundary, coincident with the DREZ, to limit Bmp signaling spreading into the intermediate spinal cord. Supporting this model, the dorsal-most dIs (dI1-dI3) are preferentially lost when netrin1 is expressed dorsally. In contrast, dP1-dP3 domains expand in the absence of netrin1. Probability of similarity between control and experimental groups: *= p < 0.05, ** p<0.005, *** p<0.0005; Student’s *t*-test. Scale bar: A-D, 50µm

We then assessed whether the gain or loss of netrin1 activity can alter the level of Bmp signaling *in vivo*. Bmps act from the roof plate (RP) at the dorsal midline to pattern the surrounding tissue^11,12^. In both chicken (Figure 6B) and mouse (open arrowheads, Figure 6A, 6F) Bmp signaling can be visualized as a graded pSmad1/5/8/ signal flanking the RP^22^. The pSmad signal can also be detected on the pial surface in the dorsal-most spinal cord, extending as far as the dorsal netrin1 boundary (closed arrowheads, Figure 6A). Electroporation of mouse netrin1 into the chicken spinal cord suppresses this signal, in a dose dependent manner. Thus, there is a >50% inhibition of pSmad activation at high (1μg) levels of netrin1 (arrows, Figure 6E, 6I), while lower (500ng, 50ng) levels of netrin1 suppress pSmad activation by 25% (arrows, Figure 6C, 6D, 6I). In contrast, we observed that the area of pSmad activation is expanded by ∼40% in spinal cords taken from *netrin1* mutant mice, compared to littermate controls (Figure 6F, 6G, 6I). Thus, the level of netrin1 has an inverse effect on the activation of Bmp signaling, consistent with the model that netrin1 acts directly as a Bmp inhibitor.

### The loss of netrin1 activity alters the level of Id signaling

Our bioinformatic analyses (Figure 5) implicated that netrin1 also regulates Bmp target genes. Since Id genes are well known targets of Bmp signaling, we assessed *Id1* and *Id3* expression in control (Figure 7A, 7C) and *netrin1^−/−^* (Figure 7B, 7D) spinal cords. Loss of netrin1 results in ∼35-40% increase in the area of both *Id1* (bracket, Figure 7E) and *Id3* (bracket, Figure 7F) expression in the dorsal most spinal cord. Taken together, these data suggest that the loss of netrin1 results in increased Bmp signaling, which in turn upregulates Id levels. Increased Id expression is predicted to maintain neural progenitors in an undifferentiated state, consistent with our observation that the loss of netrin1 results in more dPs at the expense of the dIs (Figures 4M, 4N, 7G, 7H).

**Figure 7:**
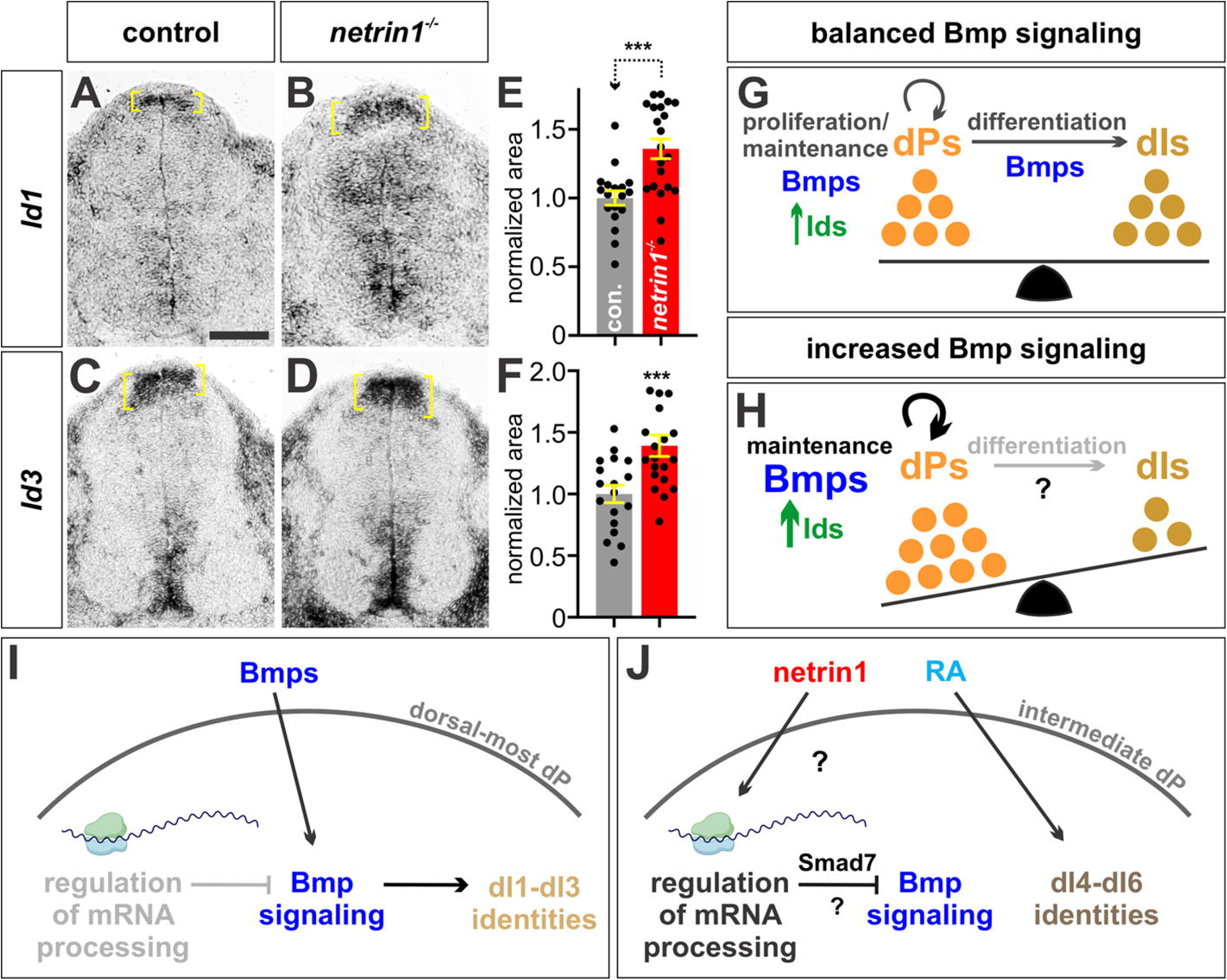
*Ids* expression is increased after the loss of *netrin1* (A-D) Control (A, C) or *netrin1^−/−^* (B, D) E11.5 mouse spinal cords were assessed for *d1* (A, B) and *Id3* (C, D) expression in the dorsal-most spinal cord (brackets). (E, F) There is a ∼35% increase in the domain of *Id1* expression (E) and a ∼40% increase in *Id3* expression (F) in the *netrin1^−/−^* dorsal-most spinal cord compared to control littermates, consistent with increased Bmp signaling (n= 20 sections, from 4 embryos). (G, H) Bmps have sequential roles in the specification of dIs, first directing dP proliferation, and then the differentiation of dPs into dIs^18^ (G). In the absence of netrin1, we observe increased Bmp and Id signaling, and an increased number of dPs, but a decreased number of dIs (H). Since Ids are a known target of Bmp signaling, these data suggest that elevating Bmp signaling directly increases Id activity, which then maintains progenitors in an undifferentiated state, and suppresses the transition to dIs. (I, J) In the dorsal most spinal cord, Bmps act from the RP to activate Bmpr signaling in NPCs, and thereby resulting the activation of Ids, and the other factors, needed for dP identity (I). In the intermediate spinal cord, the presence of netrin1 acts to limit Bmp signaling, potentially through the regulation of mRNA processing, thereby permitting intermediate dP identity (J) Probability of similarity between control and experimental groups: *** p<0.0005, Student’s *t*-test. Scale bar: 100µm

## Discussion

Netrin1 was first shown to suppress Bmp signaling in cell lines *in vitro*^41^. Here, we have used gain and loss of function approaches to examine whether netrin1 modulates Bmp signaling in the developing spinal cord using a combination of *in vivo* and *in vitro* model systems. Together, these studies suggest that netrin1 acts to limit Bmp signaling to the dorsal most spinal cord.

Previous studies have shown that Bmp signaling directs the dI1-dI3 spinal identities (Figure 7I)^11,12^, and that the extent of Bmp signaling can be inferred from a domain of pSmad1/5/8 activity extending into the VZ^22^. Our studies support the model that a further critical difference between the dorsal most dPs and the intermediate dPs is the presence of netrin1 (Figure 7I, 7J). Netrin1 has an upper boundary in intermediate dorsal spinal cord, which inversely correlates with the position of the pSmad1/5/8 domain (Figure 6A). We have found that increased netrin1 levels reduces pSmad1/5/8 activity and results in the preferential loss of dI1-dI3 (Figure 2). While the highest levels of netrin1 in the gain of function experiments appeared to generally promote cell death, we were able to identify a level of netrin1 mis-expression where we observed the specific loss of dI1-dI3, with no concomitant increase in apoptosis (Figure 2Q, 2R). In contrast, we found that the loss of netrin1 function specifically expands the pSmad1/5/8 domain, apparently resulting in an increase in the number of dP1-dP3s. These dPs appear to be maintained in a progenitor state, because the number of dI1s-dI3s concomitantly decrease (Figure 4N). Previous studies have shown that Bmps are reiteratively required during neurogenesis, having roles in progenitor maintenance, proliferation and differentiation^18,68^. Here, we observed that *Id* expression, transcriptional targets of Bmp signaling that block differentiation, was increased in the absence of netrin1 (Figure 7). Thus, the loss of netrin1 appears to elevate Bmp signaling in a manner that drives higher levels of Id expression, which then maintains dPs in a progenitor state (Figure 7G, 7H).

Taken together, we propose that the netrin1 boundary constrains the influence of Bmp signaling to the dorsal most region of the developing spinal cord (Figure 6L). The ability of netrin1 to modulate Bmp signaling in the spinal cord is a novel role for netrin1, which is best known for its role as an axon guidance cue.

### Assessing the mechanisms by which netrin1 mediate cell fate specification

The netrin1 boundary in the intermediate spinal cord is immediately adjacent to the dI1-dI3 domain (Figure 6A, 6L) suggesting that netrin1 could inhibit Bmp signaling by a direct physical interaction with Bmp ligands. A number of extracellular Bmp antagonists, including gremlin and noggin, are thought to act by sequestration, i.e., as a sink to prevent Bmp ligands from binding to the Bmp receptor^69^. We attempted multiple approaches to detect such an interaction, including computational modeling, kinetic binding assays and co-immunoprecipitation, but could find no evidence of a direct interaction between the two proteins (data not shown).

Supporting this conclusion, we found no evidence of a direct interaction between Bmp4 and netrin1 in the directed differentiation protocols. If a physical interaction between netrin1 and Bmp4 existed, then the presence of netrin1 should have inhibited Bmp4 function in all conditions tested (Figure 3A). However, we rather observed that dorsalization was only suppressed when netrin1 was added for a prolonged period of time to cells already in the dP state (condition 3). When netrin1 was added concomitantly with Bmps to cells in an earlier competence state (condition 1), there was minimal effect on transcriptional activity (Figure 5B, 5C), and no effect on dorsalization.

These bioinformatic analyses suggested an alternative mechanistic hypothesis: that netrin1 acts through the regulation of mRNA processing to suppress Bmp signaling. Our bioinformatic analyses founded that extended netrin1 treatment resulted in the upregulation of genes associated with mRNA processing and splicing. Netrin1 was previously implicated in mRNA processing in *Aplysia*, where it was found to promote local translation of ubiquitously expressed RNAs to provide spatial control during synapse formation^70^. Thus, the presence of netrin1 may stabilize the production of Bmp inhibitors known to act in the intermediate spinal cord, such as the inhibitory Smad, Smad7^71^. When Smad7 is ectopically expressed in the developing spinal cord, the intermediate fates are expanded at the expense of the dorsal most fates^72^. The local translation of Smad7, or other inhibitory factors, by netrin1 in the intermediate spinal cord would suppress Bmp signaling, and thereby permit the specification of the intermediate dI4-dI6 fates (Figure 7J).

### Autonomy vs non-autonomy of netrin1 signaling

A conundrum in these studies is that netrin1 is expressed in the intermediate spinal cord, but yet results in a non-autonomous loss of function phenotype, expanding the size of progenitor domains in the dorsal most spinal cord, coupled to the loss of dI1-dI3. We nonetheless hypothesize that netrin1 does function autonomously in the intermediate spinal cord to regulate cell fate. Netrin1 is a member of the laminin superfamily, i.e., an extracellular matrix component, that is thought to act at very short range to control axon guidance decisions^28,29^. Thus, it is very unlikely that netrin1 acts by diffusion, such that it could directly reach, and signal to, more dorsal cells. Rather, we hypothesize that removing netrin1 permits increased Bmp signaling, which results in an expansion in the size of the dorsal spinal cord. The dP1-dP3s adjust their boundaries in a compensatory manner, thereby resulting in larger domains (Figure 6L). The most profound effect is on the area of the domain, but there is also an increase in the number of dPs. As already discussed, this effect may result from the maintenance of the progenitor state, given that we do not observe an increase in proliferation (Figure 4C, 4F, 4G).

It also remains unresolved where netrin1 and Bmp signaling interact in the intermediate spinal cord. Our current model predicts that netrin1 acts on intermediate progenitor cells to prevent them from responding to the Bmp ligand. While there are low levels of netrin1 protein in the VZ, VZ-derived protein is present at highest levels on the pial surface^28,29^ (Figure 6A). Interestingly, pSmad1/5/8 is also present on the pial surface, with an inverse distribution that of pial-netrin1 (closed arrowheads, Figure 6A). However, the pial surface is not a cellular substrate, making it unlikely that this is the site of the netrin1/Bmp interaction. It may rather be a read out of proteins that are trafficked along the radial processes of the neural progenitors, again supporting the hypothesis that the domain of Bmp signaling is immediately adjacent to the netrin1 domain.

### Role of canonical netrin1 receptors mediating suppression of Bmp signaling

Netrin1 binds to a number of different receptors, including Dcc, neogenin1 and members of the Unc5 family, many of which are present in the dorsal spinal cord^29,73,74^. It remains unresolved which of these receptors mediates the ability of netrin1 to regulate Bmp signaling. Our previous studies have shown that *Dcc* is consistently expressed at low levels in the dorsal and intermediate VZ, while *neogenin1* has a more dynamic expression pattern first in the intermediate VZ and then in the dorsal most progenitors^74^. The studies using the stem cell directed differentiation model identified a putative time window when the netrin1 cell fate receptor(s) might function. We observed that netrin1 treatment only affects the transcriptional status of stem cell derived progenitors when added on day 5, but not at day 4 (Figure 5B, 5C). One explanation for this observation is that the receptor complement needed to respond to netrin1 is only present on day 5, i.e., once spinal progenitors transition to the dP state. Analysis of the scRNA-Seq atlas derived from the RA±Bmp4 protocols^57^ (https://samjbutler.shinyapps.io/Data_Viewer/) shows that Dcc, neogenin1, Unc5c and Unc5d are expressed at different timepoints along the differentiation trajectory (Figure S1B-S1E). However, only Dcc has the profile that fits the observed netrin1 responsiveness, i.e., that the expression of Dcc is upregulated immediately after transitioning from the progenitor state.

While this analysis supports the hypothesis that Dcc is a cell fate receptor, other receptors must also be required. Dcc is not present in the chicken genome, and neogenin1 has been proposed as the functional substitute^74^. Netrin1 was able to suppress pSmad1/5/8 activity in a dose dependent manner in Cos7 cells after Bmp4 stimulation (Figure 6G, K). However, we were unable to detect that either Dcc or neogenin1 are present in Cos7 cells by western analyses (data not shown). Future studies will assess whether netrin1 is interacting Dcc and/or a novel receptor to inhibit Bmp signaling.

### Broader implications

The Bmp family of growth factors is used to specify developmental decisions in all organ systems, in a manner that is conserved across species. While netrin1 was first studied for its axon guidance activities in the nervous system, subsequent studies have shown that it plays critical roles in the development of other organs, including the kidney, lungs, and mammary glands, as well as in the progression of diseases, such as cancer and diabetes. Thus, it is likely that the interaction between netrin1 and Bmp signaling observed in these studies will be critical for other developmental and disease processes, potentially as fine-tuning mechanism that permits topographic regulation.

## Authors Contributions

SA and SJB conceived the project. SA and KH analyzed the distribution of netrin1 and netrin2. SA and YMA performed the chicken gain of function analyses, while SA, YMA and SG performed the mouse loss of function analyses. SA assessed the Bmp/netrin1 interaction in vitro. SG performed the cellular differentiations, and conducted bulk RNA Seq. SG and RK analyzed the bulk RNA-Seq data. This paper was written by SA, SG and SB and edited by SA, SG and SJB.

## Acknowledgements

We would like to thank Lisa Goodrich, Thomas Muller, Bennett Novitch and Marc Tessier-Lavigne for mouse lines and reagents, and Avihu Klar, Bennett Novitch Alvaro Sagasti, Larry Zipursky and the members of the Butler and Novitch laboratories for discussions. One illustration was made using our institutional site license from Biorender.com This work was supported by a UCLA senior undergraduate research scholarship to K.H.; graduate fellowships from the NIH (F31 GM007185, T32 GM7185) and UCLA (Eugene V. Cota-Robles, Whitcome and Hilliard Neurobiology award) to S.A. and NSF (DGE-2034835) to Y.M-A.; a UCLA Broad Stem Cell Research Center (BSCRC) postdoctoral training grant (to S.G); and grants from the National Institutes of Health (NIH) (P50 HD103557) to R.K and (R01 NS123187, R01 NS085097, P50 HD103557) and innovation awards from the BSCRC to S.J.B.

## Inclusion and Diversity

One or more of the authors of this paper self-identifies as an underrepresented group in science. One or more of the authors of this paper self-identifies as a member of the LGBTQ+ community. While citing references scientifically relevant for this work, we also actively worked to promote gender balance in our reference list.

## STAR Methods

### RESOURCE AVAILABILITY

#### Lead Contact

Further information and requests for resources and reagents should be directed to Samantha Butler.

#### Materials availability

Information regarding the resources and reagents used in this paper should be directed to the lead contact, Samantha Butler.

#### Data and Code Availability

Bulk RNA-seq data has been deposited at GEO and is publicly available. Accession numbers are listed in the key resources table.

Microscopy data reported in this paper will be shared by the lead contact upon request. Any additional information needed to reanalyze the data reported in this paper is available from the lead contact upon request.

### KEY RESOURCES TABLE

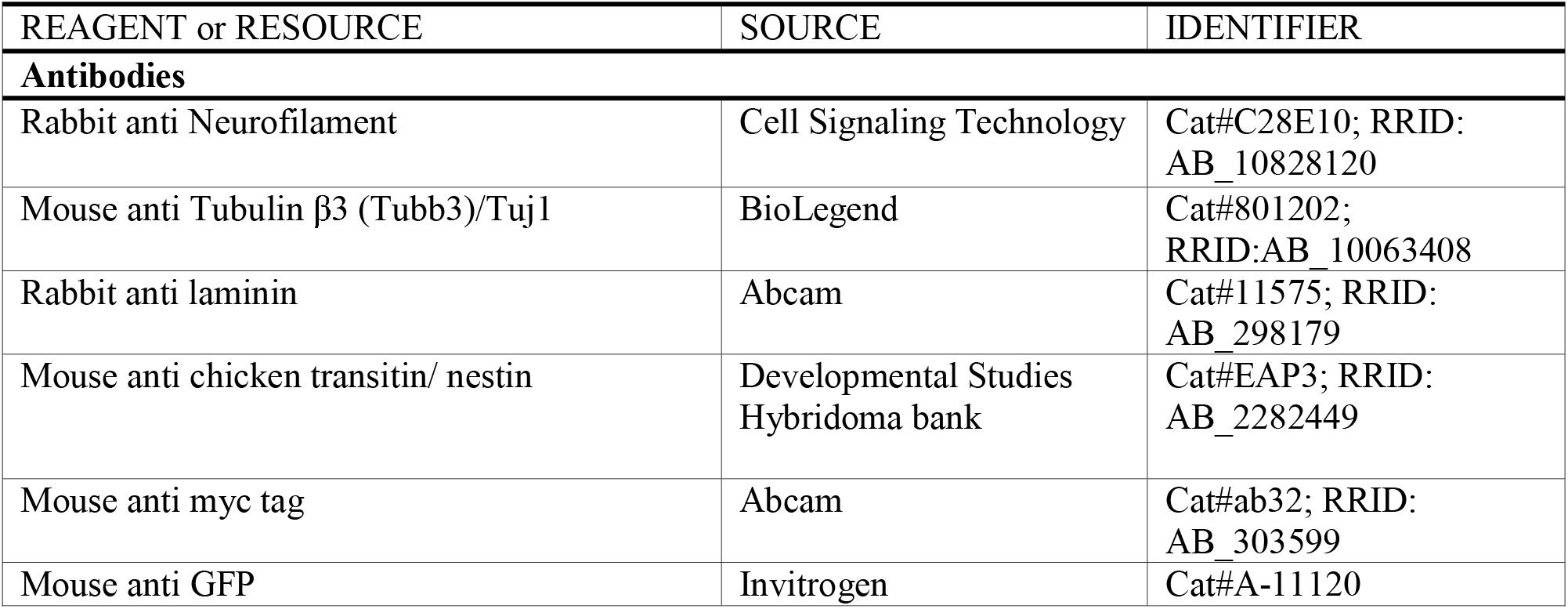
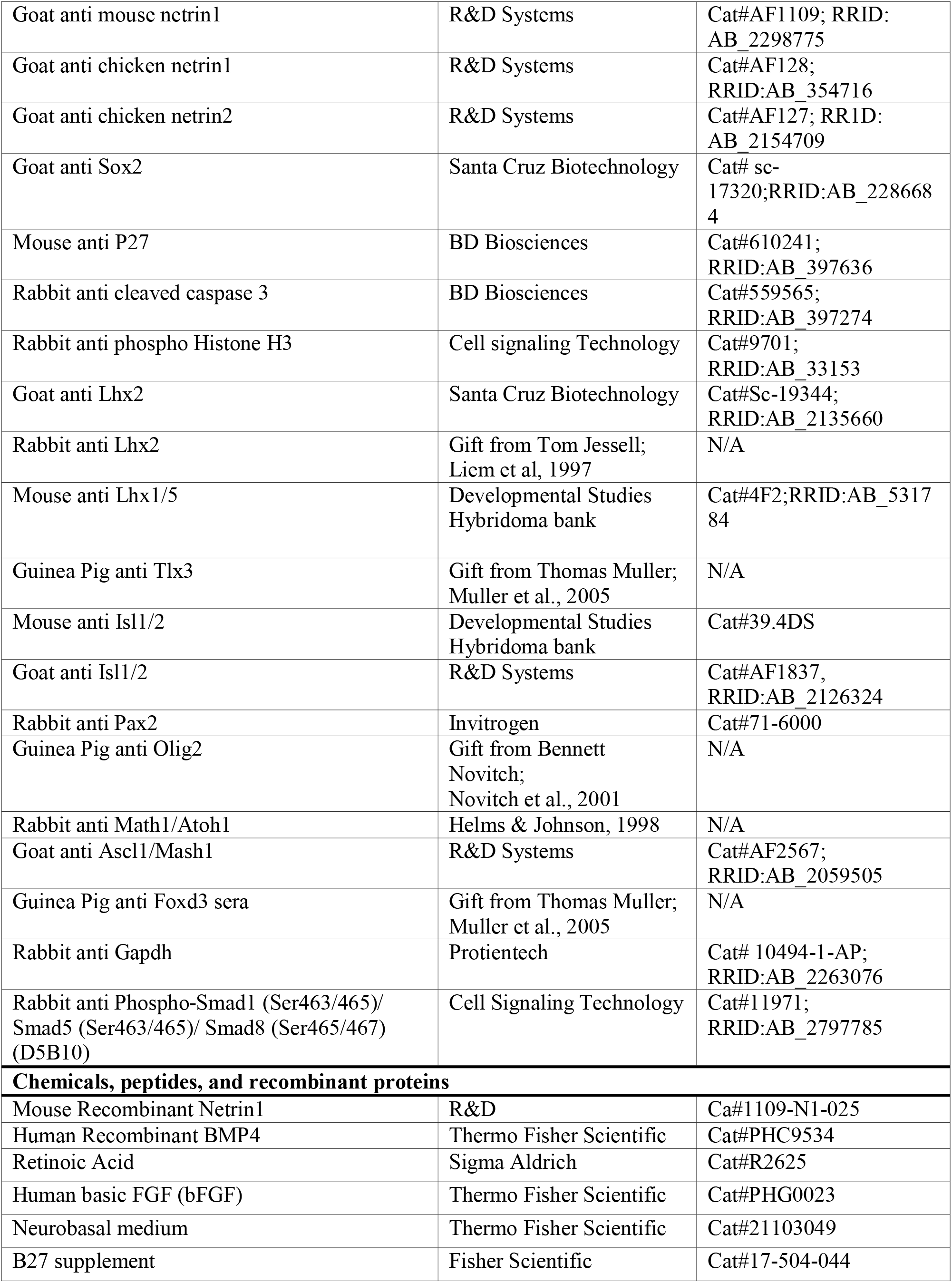
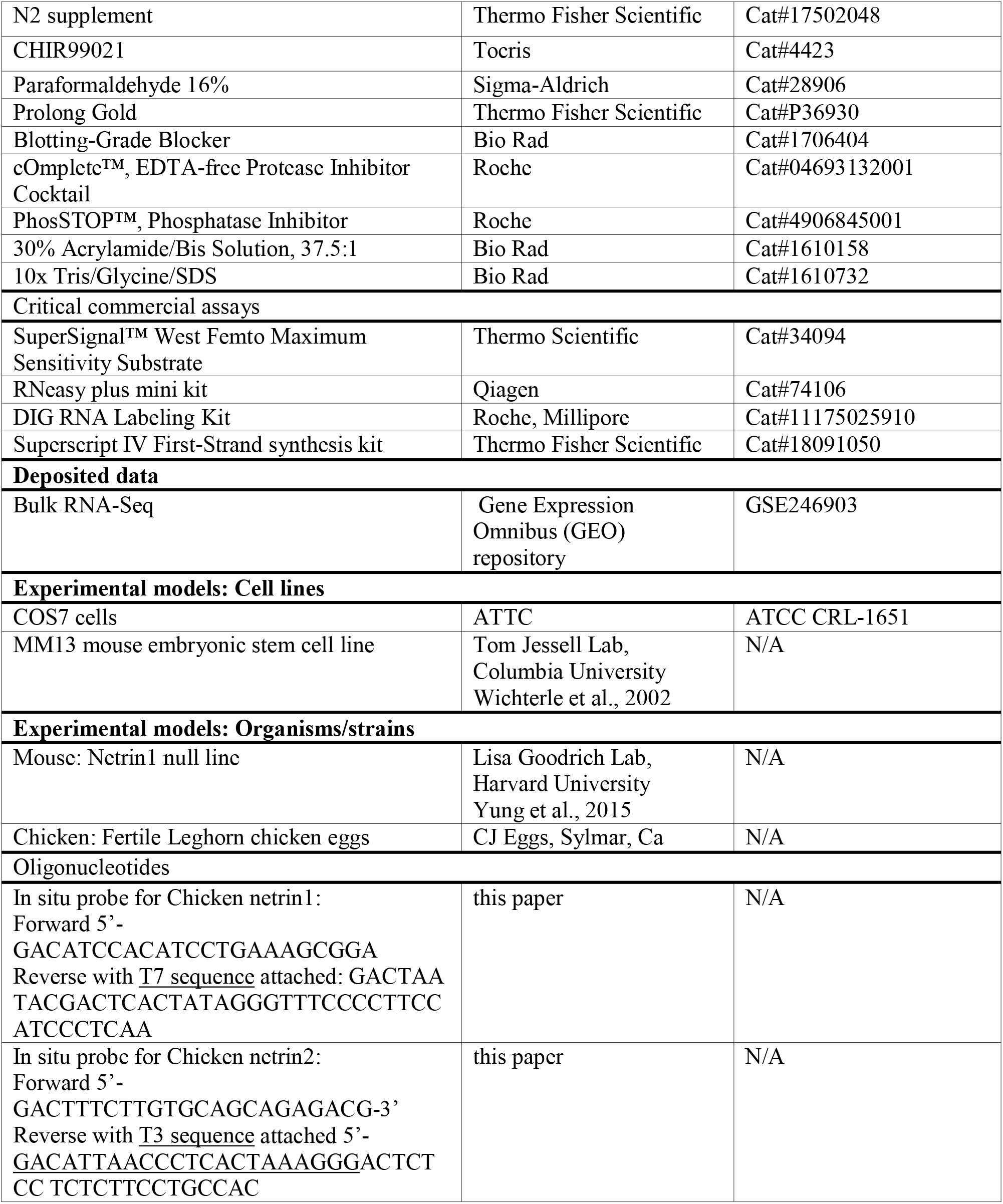
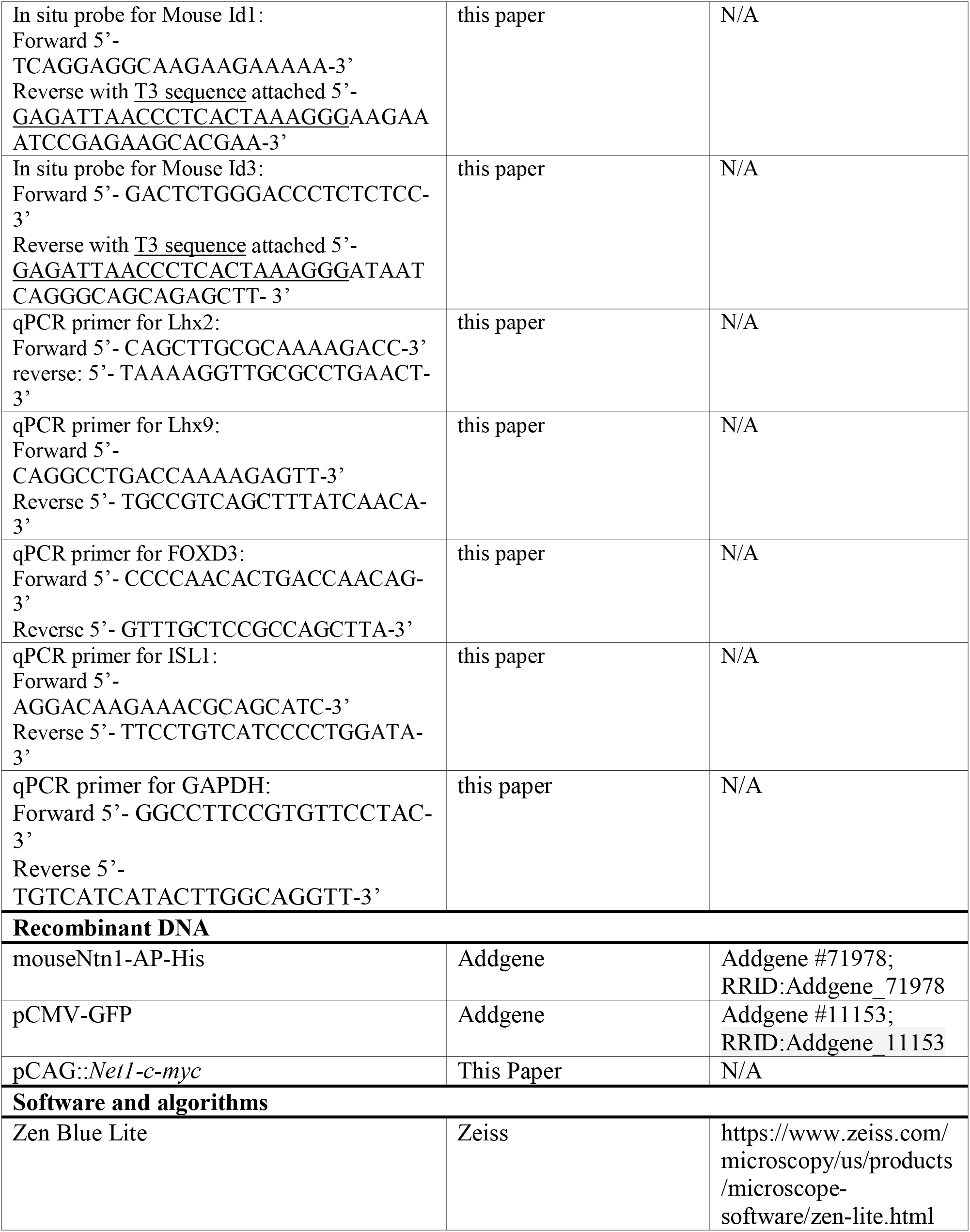
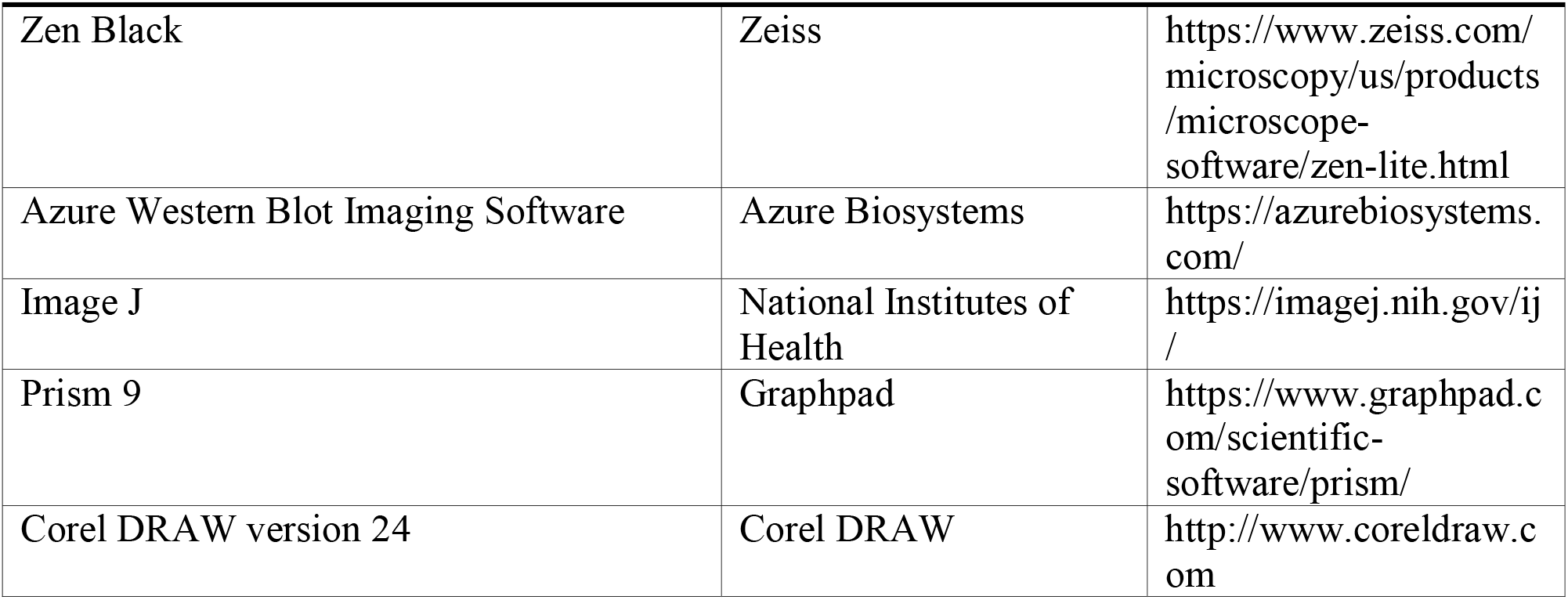

### EXPERIMENTAL MODEL AND STUDY PARTICIPANT DETAILS

#### Chicken Embryos, Mice Generation, and Culture of Cos7 cell lines

Fertile Leghorn eggs (CJ Eggs, Sylmar CA) were incubated for 60 hours until the embryos reached Hamburger Hamilton stages 14–15. The spinal cord was *in ovo* electroporated as previously described^20^ and allowed to develop until HH stages 24–26, at which time the tissue was analyzed.

The netrin1 null line was a gift from Dr. Lisa Goodrich^60^. Embryos were collected from timed matings, and the presence of a vaginal plug was considered embryonic day (E) 0.5. Heads were used to isolate the DNA and were amplified by PCR to identify the genotypes of each embryo^60^. All animal procedures were carried out in accordance with University of California Los Angeles IACUC guidelines.

Cos7 cells (ATCC CRL-1651) were cultured in Dulbecco’s modified Eagle’s medium (DMEM) (Sigma-Aldrich) supplemented with 10% fetal bovine serum (FBS) (Fisher Scientific) and Penicillin-Streptomycin-Glutamine (100X) (Gibco, Fisher Scientific).

### METHOD DETAILS

#### In ovo electroporation of chicken embryos

A c-myc tag (EQKLISEEDL) was fused to the C terminal end of mouse *netrin1* (Addgene #71978) using PCR cloning. Netrin1-myc was then subcloned into the CAGGS vector, under the control of the CAG enhancer^75^, which is comprised of a CMV enhancer and chicken β-actin promoter. Fertile Leghorn eggs (CJ Eggs, Sylmar CA) were incubated for 60 hours until the embryos reached Hamburger Hamilton stages 14–15. The spinal cord was *in ovo* electroporated as previously described^20^ and allowed to develop until HH stages 24–26, at which time the tissue was analyzed.

The following constructs were used: CAG:*gfp* (1000ng/μl), CAG::*ntn1-myc* (50ng/μl, 500ng/μl, or 1000ng/μl). In all cases, the presence of GFP demonstrates electroporation efficiency. When altering the concentration of *netrin* expression the CAG::*ntn1-myc* expression vector was diluted with the pCAGEN vector (plasmid #11160, Addgene), to hold the concentration of DNA constant at 2000ng/μl across all experiments.

#### Tissue processing

Spinal cords were fixed using 4% paraformaldehyde for 2 hours at 4 °C. After fixation, the tissue was cryoprotected in a 30% sucrose solution overnight, after which the tissue was mounted in optimal cutting temperature (OCT) and cryosectioned at 20µm. Sections were collected on slides and processed for immunohistochemistry.

#### Immunohistochemistry

Chicken embryonic spinal cords and mouse embryonic spinal cords were sectioned to yield 20μm sections. The details of the antibodies used for immunostaining can be found in the Key Resources Table. Species appropriate Cyanine 3, 5 and Alexa Fluor 488 conjugated secondary antibodies were used (Jackson ImmunoResearch Laboratories). Images were collected on Carl Zeiss LSM700 and LSM800 confocal microscopes.

#### In situ hybridization

*In situ* hybridizations were performed on chicken (HH stage 18–25) and mouse (E11.5) embryonic spinal cords. 3’UTR probes were designed using http://primer3plus.com and verified for specificity to the gene of interest using http://www.ncbi.nlm.nih.gov/tools/primer-blast/.

The chicken and mouse primer sequences used to make *in situ* hybridization probes can be found in Key Resources Table. Probes were made using a DIG RNA labeling kit (Roche, Cat#11175025910). Images were collected on a Carl Zeiss AxioImager M2 fluorescence microscope with an Apotome attachment.

#### Western blot analyses

Cos7 cells were seeded in 12-well or 24 well plates the day before stimulation with netrin1 (R&D, cat# 1109-N1-025) and Bmp4 (Thermo Fisher cat# PHC9534). On the day of stimulation, the cells were starved in FBS-free media for an hour prior to stimulation with 5ng/ml Bmp4and 0.5μg/ml, 0.25 μg/ml or 0.125 μg/ml netrin1. After one hour of stimulation, the cells were washed with PBS and lysed with RIPA lysis buffer containing a protease inhibitor cocktail (Roche) and the phosphatase inhibitor PhosSTOP (Roche). The cell lysates were kept on ice for 30 min and centrifuged at 20,800×g for 10 min at 4 °C. The supernatant was subjected to SDS-PAGE using 10% Tris-Glycine SDS gels followed by transfer onto PVDF membranes (Millipore Sigma). The membranes were blocked using non-fat dry milk (Bio-Rad Laboratories). The blocked membranes were then incubated with the primary antibodies (Key Resources Table) at 4° C overnight. Thereafter, the membranes were washed three times with TBST (20 mM Tris, 150 mM NaCl, 0.1% Tween 20), incubated with species specific horseradish peroxidase conjugated secondary antibodies (Jackson ImmunoResearch Laboratories) for an hour at room temperature and then washed again three times with TBST. The chemiluminescent bands were analyzed using the Pierce Femto Chemiluminescence system on the Azure Imager.

#### Mouse embryonic stem cell (mESC) culture and differentiation

The mouse ESC line MM13^54^ was maintained in ESC medium with LIF on mitotically inactive MEFs. Before differentiation, ESC colonies were dissociated, plated on gelatin-coated plates, and allowed to proliferate for 1–2 days. To initiate differentiation, cells were plated on 0.1% gelatin-coated 24-well CellBIND dishes (Corning) with N2/B27 medium containing 10ng/ml basic fibroblast growth factor (bFGF). On day 1, small colonies of cells can be observed attached to the bottom of the wells. On day 2, cells were supplemented with 10ng/ml bFGF and 5μM CHIR99021 (Tocris, Cat#4423) for 24 h to induce a neuromesodermal identity^55^. On day 3, cells were directed towards a spinal lineage by exposing them to 100nM RA (Sigma Aldrich, cat# R2625) for 24 hours, followed by 100nM RA + 10ng/ml Bmp4 (Thermo Fisher cat# PHC9534) to induce dorsal spinal cord identity^25^. To evaluate the effects of netrin1 on dI differentiation, two concentrations of mouse recombinant netrin1 (high - 0.5 µg/ml, low - 0.125 µg/ml) (R&D, Cat#1109-N1-025) were added in three different timelines (conditions 1, 2, 3).

For condition 1, netrin1 was added with RA + Bmp4 between day 4 and day 5, resulting in 24 hours of netrin1 exposure. For condition 2, netrin1 was added between day 5 and day 6, providing 24 hours of netrin1 exposure after the initial patterning by RA+Bmp4. For condition 3, netrin1 was added every other day between day 5 to day 9, leading to an extended 4-day netrin1 exposure. Terminal differentiation was induced by replacing the growth factor containing media with basic N2/B27 medium at day 5, and cultures were allowed to differentiate until day 9. At the end of the differentiation, the cultures were lysed in buffer RLT and RNA was purified using RNAeasy kit (Qiagen, Cat#74104) for preparing cDNA for quantitative reverse transcriptase PCR analysis.

#### Reverse transcriptase PCR analysis

RNA was extracted from at least two independent differentiations using the RNeasy mini purification kit (Qiagen, Cat#74104). cDNA was synthesized using Superscript IV (Thermo Fisher Scientific, Cat#18091050) using oligodT as primers to convert mRNAs into cDNAs. RT-qPCR was always performed in triplicate using SYBR Green Master Mix on a Roche RT-qPCR thermocycler using gene-specific primers (Key Resources Table). The Ct values for each gene were calculated by averaging three technical replicates from independent differentiations for each condition. Expression of the target gene was normalized with the expression of glyceraldehyde-3-phosphaste dehydrogenase (GAPDH) and fold change was calculated using the 2^−ΔΔCt^ method^76^. The variation in fold change in expression is represented in ±SEM (standard error of mean).

#### Bulk RNA-Seq and bioinformatic analysis of mESC differentiation cultures treated with recombinant netrin1

RNA samples for bulk RNA-Seq were collected on day 5 (condition 1), day 6 (condition 2) and day 9 (condition 3) using an RNAeasy mini kit (Qiagen). Libraries were prepared using universal plus mRNA sequencing kit (NuGEN) and sequenced onto Novaseq S2 to obtain minimum 30-50 million reads/samples. Reads were then aligned to latest mouse genome assembly (mm10) using the STAR spliced read aligner and differentially expressed genes were obtained using EdgeR^77^ and DEseq2 packages on R with a selection criteria of FDR p<0.05. The differentially expressed genes were subjected to Gene Ontology (GO) analysis using Enrichr^78^. The top 10 GO categories were selected for demonstration in figure 5. For heatmaps, genes that fall under a specific GO category (e.g., mRNA processing) were identified and their FPKM values were extracted and plotted using Heatmapper.ca with row scaling.

### QUANTIFICATION AND STATISTICAL ANALYSIS

#### Quantification

For the electroporation experiments, the non-electroporated side of the spinal cord was used as the negative control in cell-counting experiments. Thus, the cell number on the electroporated side was normalized against a GFP control electroporation performed at the same time. For the quantification of pSmad1/5/8 intensity, control and experimental embryos were stained on the same slides. The staining intensity and area were quantified using the ImageJ software. Biological replicates: 1–2 chicken embryos per experimental condition were collected within an experiment. Each experiment was repeated at least three times. Technical replicates:>10 or more sections per embryo were analyzed.

For the transgenic mice experiments, cell counts, area and stain intensities were normalized to the littermate controls. Biological replicates: 1-2 embryos per experimental condition were collected within an experiment, and embryos were collected from at least three timed pregnancies (for a total of 4-6 embryos/condition. All analyses were performed using littermate controls. Cell counts were performed in manner blinded to genotype. Technical replicates:>10 or more sections per embryo were analyzed.

For cell culture experiments, experiments were performed at least 3 independent times. All values were normalized to internal controls.

#### Statistics

Data are represented as mean ± SEM (standard error of the mean). Tests for statistical significance were performed using Prism software (version 9). Values of p < 0.05 were considered significant in all cases.

