## Supplemental figures for "Netrin1 patterns the dorsal spinal cord through modulation of Bmp signaling"

Supplemental Figures and Tables

**Supplemental Figure 1:** Expression of netrin1 and key receptors in directed differentiated protocols

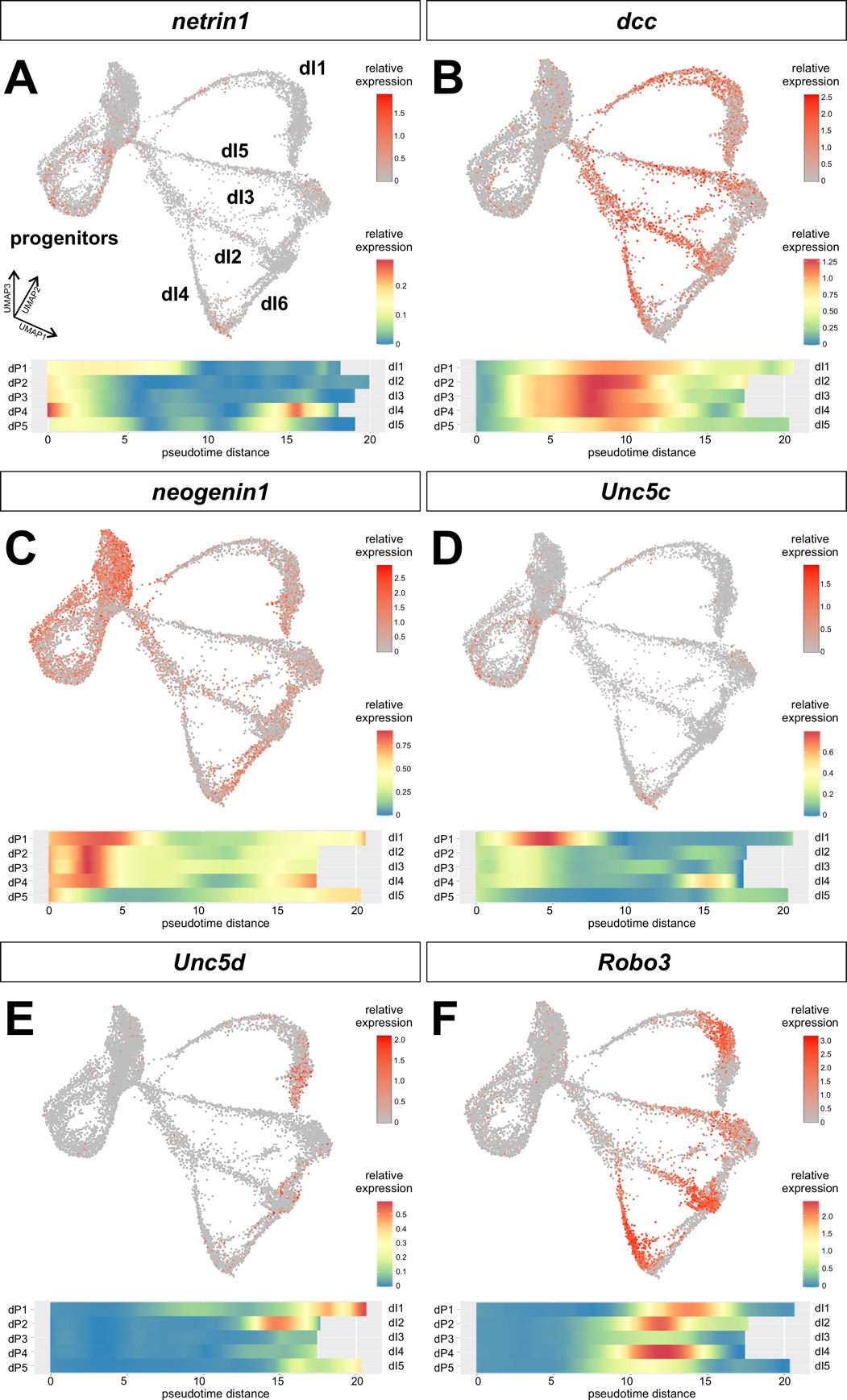

Single cell RNA atlas of the RA±BMP4 directed differentiation protocols^57^, annotated with the major lineage derivatives (A) The pseudotime values from the atlas show r timeline of expression after the progenitor state. The entire data set can be viewed here: <https://samjbutler.shinyapps.io/Data_Viewer/> .

(A-F) *Netrin1* is expressed at highest levels in a subset of cycling progenitors (A). *Dcc* is specifically upregulated in dPs and dIs (B). *Neogenin1* is present at highest levels in progenitors, and lower levels in the dPs and dIs (C). *Unc5c* is expressed at highest levels in a subset of cycling progenitors (D). *Unc5d* is present in highest levels in differentiated dI1s and dI2s (E). *Robo3* is present at high levels in dIs as they are maturing, but not in their most mature states (F).

**Supplemental Figure 2:** Netrin1 overexpression results in the loss of commissural axons in the developing chicken spinal cord.

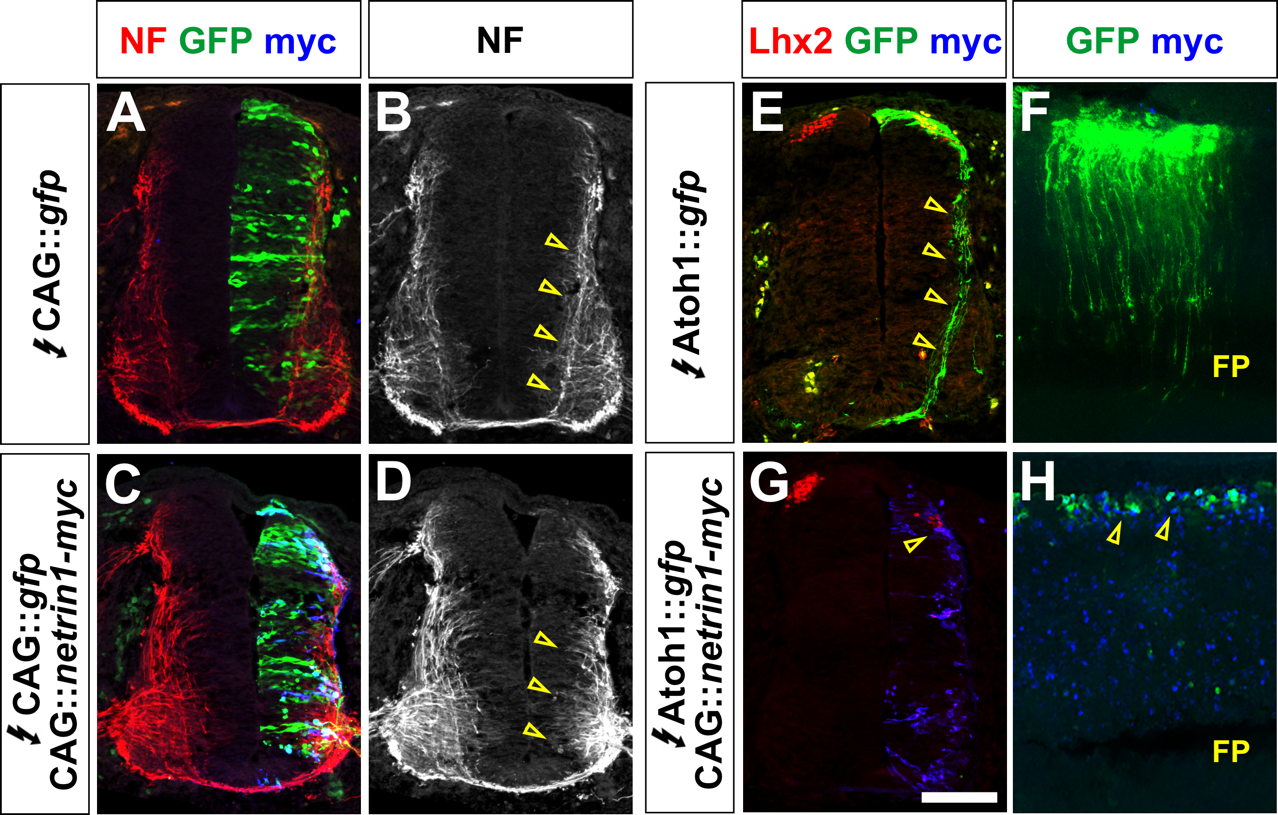

(A-H) Chicken spinal cords were electroporated at HH stage 14 with *Gfp* under the control of the CAG (A-D) or Atoh1 (E-H) with alone (A, B, E, F) or with CAG::*netrin1* (C, D, G, H) and incubated until HH stage 24. Thoracic transverse sections (A-D, E,.G) or longitudinal whole mounts (F, H) were labeled with antibodies against neurofilament NF, (red, A-D), Lhx2 (red, E, F), GFP (green, A, C, E-H) and myc (blue, A, C, E-H)

(A-D) In control embryos, NF^+^ axons are present in both the motor column, and projecting circumferentially around the VZ (arrows, B). These commissural axons are not present in *netrin1* electroporated embryos (arrows, D),

(E- H) In control embryos, the Atohl1 promoter drives the expression of GFP specifically in the commissural axons of the dI1 population (arrows, E), which project to the floor plate (FP). However, dI1 axons are not observed in the transverse sections of *netrin1* electroporated embryos. The longitudinal view reveals some electroporation in cells at the dorsal most edge, but confirms the lack of any dI1 axons.

**Supplementary Table 1:** Regulatory transcription factors associated with the downregulated genes in condition 3.

| **Regulatory TF** | **P value** | **Genes downregulated in condition 3** |
| --- | --- | --- |
| Aatf | -1.757267094 | TRP53, BAX, TPT1, BBC3 |
| Trp73 | -1.612517391 | PDGFRB, PSRC1, TRP53, E2F1, BAX, FOXO3, TPT1, BBC3 |
| Myc | -1.561997835 | PDGFRB, NOP56, TRRAP, CDKN2A, CAD, CDCA7, NRGN, ZFP36, MTA1, TERT, CDK4, TRP53, SNAI1, ACP5, BAX, WNT4 |
| Foxm1  (human) | -1.406064866 | FGB, BTG2, CDKN2A, VEGFB, CDC6, APOE, CDC25A |
| Trp53 | -1.32745725 | BTG2, MCM7, SRC, UHRF1, PSEN1, SLC2A4, AFP, FOXO3, FOXM1, PKD1, RELA, BBC3, SALL2, TERT, ZFP385A, RPS6KA1, E2F1, HMOX1, E4F1, GTSE1, HRAS, PTPRV |
| Nrf1 | -1.198846867 | PRDX3, SLC46A1, PRDX5, FXR2, CAPNS1, COX6A1 |
| **Egr1** | -1.176264929 | CDKN2D, **JUND**, PTGES2, **ENO1**, **MAPK14**, CACNA1H, ICAM1, **SMAD7**, RCAN1, MAP1LC3B, MMP14, **SOCS1**, STIM1, TRP53, PP1R1B, **ID3**, BAX, TK1, PTGES, **WNT4** |
| Foxm1  (mouse) | -1.127701227 | SOCS3, MYCN, PTTG1, CDKN2A, LIF, SNAI1, PRRC2A, NFKB2 |
| Tp53 | -1.117276528 | BTG2, MCM7, UHRF1, OGG1,GSTP1, SLC2A1, AFP, FOXO3, FOXM1, RELA, BBC3, PTPA, RECQL4, RASSF1, PTTG1,TERT, POLD1, HSF1, FDXR, E2F1, AKT1, CTSD, HRAS |
